# SARS-CoV-2 proteins exploit host’s genetic and epigenetic mediators for the annexation of key host signaling pathways that confers its immune evasion and disease pathophysiology

**DOI:** 10.1101/2020.05.06.050260

**Authors:** Md. Abdullah-Al-Kamran Khan, Abul Bashar Mir Md. Khademul Islam

**Author notes:** **Correspondence:** Dr. Abul Bashar Mir Md. Khademul Islam, Associate Professor, Department of Genetic Engineering and Biotechnology, University of Dhaka, Dhaka 1000, Bangladesh.

## Abstract

The constant rise of the death toll and cases of COVID-19 has made this pandemic a serious threat to human civilization. Understanding of host-SARS-CoV-2 interaction in viral pathogenesis is still in its infancy. In this study we aimed to correlate how SARS-CoV-2 utilizes its proteins for tackling the host immune response; parallelly, how host epigenetic factors might play a role in this pathogenesis was also investigated. We have utilized a blend of computational and knowledgebase approach to elucidate the interplay between host and SARS-CoV-2. Integrating the experimentally validated host interactome proteins and differentially expressed host genes due to SARS-CoV-2 infection, we have taken a blend of computational and knowledgebase approach to delineate the interplay between host and SARS-CoV-2 in various signaling pathways. Also, we have shown how host epigenetic factors are involved in the deregulation of gene expression. Strikingly, we have found that several transcription factors and other epigenetic factors can modulate some immune signaling pathways, helping both host and virus. We have identified miRNA hsa-miR-429 whose transcription factor was also upregulated and targets were downregulated and this miRNA can have pivotal role in suppression of host immune responses. While searching for the pathways in which viral proteins interact with host proteins, we have found pathways like-HIF-1 signaling, autophagy, RIG-I signaling, Toll-like receptor signaling, Fatty acid oxidation/degradation, Il-17 signaling etc significantly associated. We observed that these pathways can be either hijacked or suppressed by the viral proteins, leading to the improved viral survival and life-cycle. Moreover, pathways like-Relaxin signaling in lungs suggests aberration by the viral proteins might lead to the lung pathophysiology found in COVID-19. Also, enrichment analyses suggest that deregulated genes in SARS-CoV-2 infection are involved in heart development, kidney development, AGE-RAGE signaling pathway in diabetic complications etc. might suggest why patients with comorbidities are becoming more prone to SARS-CoV-2 infection. Our results suggest that SARS-CoV-2 integrates its proteins in different immune signaling pathway and other cellular signaling pathways for developing efficient immune evasion mechanisms, while leading the host to more complicated disease condition. Our findings would help in designing more targeted therapeutic interventions against SARS-CoV-2.

## 1. Introduction

Though several human coronaviruses outbreaks caused severe public health crisis over the past few decades, the recent coronavirus disease (COVID-19) outbreak caused by the Severe Acute Respiratory Syndrome Coronavirus 2 (SARS-CoV-2) has beaten the records of the previous and still the case counts are still on the upswing. About 210 countries and territories around the globe has been affected by this outbreak and around a total of 2 millions of people are already infected with SARS-CoV-2 and the number is steadily rising till the date of writing this article (Worldometer, 2020). Out of the closed cases, almost 20% of the patients have suffered death and about 5% of the active cases are in critical conditions (Worldometer, 2020). Though the death rates from COVID-19 was estimated to be as small as 3.4% (WHO, 2020), at present the global fatality rate is changing very rapidly; therefore, more comprehensive studies needto be done for the efficient controlling to overturn this pandemic. Coronaviruses are single stranded positive sense, enveloped RNA virus having ~30Kb genome (Lu et al., 2020). Amongst the four genera, SARS-CoV-2 (Accession no. NC_045512.2) belong to the betacoronavirus genus and it has ~29.9Kb genome encoding 11 genes (NCBI-Gene, 2020). SARS-CoV-2 shares about 90% genome sequence similarity with bat derived SARS-like coronavirus, whereas this novel virus have only ~79% and ~50% similarities with Severe Acute Respiratory Syndrome Coronavirus (SARS-CoV) and Middle East Respiratory Syndrome-related Coronavirus (MERS-CoV), respectively (Jiang et al., 2020; Lu et al., 2020; Ren et al., 2020). A substantial genomic difference can be observed between SARS-CoV and SARS-CoV-2; as in SARS-CoV-2, there have been 380 amino acids substitution, deletion of ORF8a, elongation of ORF8b, and truncation of ORF3b observed (Lu et al., 2020).

Though the overall mortality rate from SARS-CoV is higher than that of SARS-CoV-2, several unique features of SARS-CoV-2 like-increased incubation period and dormancy inside the host, thus spreading more efficiently (Lauer et al., 2020). This suggests that SARS-CoV-2 might be using some immune evasion strategies to maintain its survival and essential functions within its host.

Upon viral infection, host innate immune system detects the virion particles and elicits the first sets of antiviral responses (Katze et al., 2008) to eliminate the viral threats. However, viruses itself have generated various modes of actions to evade those immune response by modulating the host’s intracellular signaling pathways (Kikkert, 2020). This arm-wrestling between the host and the infecting virus results in the immunopathogenesis. Different human coronaviruses also show similar features of host-pathogen interactions, which ranges from the viral entry, replication, transcription, translation, assembly to the evasion from host innate immune response (Fung and Liu, 2019). Not only this but also different antiviral cellular responses like-autophagy (Ahmad et al., 2018), apoptosis (Barber, 2001) etc. can also be moderated by the virus to ensure its survival inside the host cells. Apart from these, several other host-virus interactions are also observed like-modulation of the activity of host transcription factors (Lyles, 2000), host epigenetic factors (e.g. histone modifications, host miRNAs etc.) (Adhya and Basu, 2010). All of these multifaceted interactions can lead to the ultimate pathogenesis and progression of the disease.

The interplay between different human coronaviruses and host was previously reported (Fung and Liu, 2019), however, SARS-CoV-2 interactions with the host immune response and its outcome in the pathogenesis are still to be elucidated. Gordon et al. (2020) identified 332 high confidence interactions between SARS-CoV-2 proteins and human proteins (Gordon et al., 2020), and Blanco-Melo et al. (2020) produced a transcriptional signatures of SARS-CoV-2 infected cells (Blanco-Melo et al., 2020). In this study, we aimed to correlate the complex host - SARS-CoV-2 interactions with the associated differentially expressed genes found in the SARS-CoV-2 infection. Also, we have compared the differential gene expression profiles of SARS-CoV and SARS-CoV-2 infected cells to find out those pathways uniquely targeted by SARS-CoV-2. Moreover, we have also incorporated other associated host epigenetic factors which might play a role in the pathogenesis by deregulating the signaling pathways.

## 2. Materials and Methods

### 2.1. Retrieval of the host proteins that interact with SARS-CoV-2

We have obtained the 332 human proteins that forms high confidence interactions with SARS-CoV-2 proteins from the study conducted previously by Gordon et al. (2020) and processed their provided proteins name into the associated HGNC official gene symbol (Supplementary file 1).

### 2.2. Analysis of microarray expression data

Microarray expression data on SARS-CoV infected 2B4 cells or uninfected controls for 24 hrs obtained from Gene Expression Omnibus (GEO), accession: GSE17400 (https://www.ncbi.nlm.nih.gov/geo) (Barrett et al., 2012). Raw Affymatrix CEL files were background corrected, normalized using Bioconductor package “affy v1.28.1” using ‘rma’ algorithm. Quality of microarray experiment (data not shown) was verified by Bioconductor package “arrayQualityMetrics v3.2.4” (Kauffmann et al., 2009). Differentially expressed (DE) between two experimental conditions were called using Bioconductor package Limma (Smyth, 2005). Probe annotations were converted to genes using in-house python script basing the Ensembl gene model (Biomart 99) (Flicek et al., 2007). The highest absolute expression value was considered for the probes that were annotated to the same gene. We have considered the genes to be differentially expressed, which have FDR (Benjamini and Hochberg, 1995) p-value ≤ 0.05 and Log2 fold change value ≥ 0.25.

### 2.3. Analysis of RNA-seq expression data

Illumina sequenced RNA-seq raw FastQ reads were extracted from GEO database accession: GSE147507 (Barrett et al., 2012). This data include independent biological triplicates of primary human lung epithelium (NHBE) which were mock treated or infected with SARS-CoV-2 for 24hrs. Mapping of reads was done with TopHat (tophat v2.1.1 with Bowtie v2.4.1) (Trapnell et al., 2009). Short reads were uniquely aligned allowing at best two mismatches to the human reference genome from (GRCh38) as downloaded from USCS database (Lander et al., 2001). Sequence matched exactly more than one place with equally quality were discarded to avoid bias (Hansen et al., 2010). The reads that were not mapped to the genome were utilized to map against the transcriptome (junctions mapping). EnsEMBLgene model (Hubbard et al., 2007) (version 99, as extracted from UCSC) was used for this process. After mapping, we used SubRead package featureCount v2.21 (Liao et al., 2013) to calculate absolute read abundance (read count, rc) for each transcript/gene associated to the Ensembl genes. For differential expression (DE) analysis we used DESeq2 v1.26.0 with R v3.6.2 (2019-07-05) (Anders and Huber, 2010) that uses a model based on the negative binomial distribution. To avoid false positive, we considered only those transcripts where at least 10 reads are annotated in at least one of the samples used in this study.

### 2.4. Functional enrichment analysis

We utilized Gitools v1.8.4 for enrichment analysis and heatmap generation (Perez-Llamas and Lopez-Bigas, 2011). We have utilized the Gene Ontology Biological Processes (GOBP) and Molecular Function (GOMF) modules (Ashburner et al., 2000), KEGG Pathways (Kanehisa and Goto, 2000), Bioplanet pathways (Huang et al., 2019b) modules and Wikipathways (Slenter et al., 2017) modules for the overrepresentation analysis. Resulting p-values were adjusted for multiple testing using the Benjamin and Hochberg’s method of False Discovery Rate (FDR) (Benjamini and Hochberg, 1995). We have also performed the enrichment analysis based on KEGG pathway module of STRING database (Szklarczyk et al., 2019) for the 332 proteins (Supplementary file 1) retrieved from the analysis of Gordon et al. (2020) (Gordon et al., 2020) along with the deregulated genes analyzed from SARS-CoV-2 infected cell’s RNA-seq expression data.

### 2.5. Obtaining the transcription factors binds promoter regions

We have obtained the transcription factors (TFs) which bind to the given promoters from Cistrome data browser (Zheng et al., 2018) that provides TFs from experimental ChIP-seq data. We utilized “Toolkit for CistromeDB”, uploaded the 5Kb upstream promoter with 1Kb downstream from transcription start site (TSS) BED file of the deregulated genes and fixed the peak number parameter to “All peaks in each sample”.

### 2.6. Obtaining human miRNAs target genes

We extracted the experimentally validated target genes of human miRNAs from miRTarBase database (Huang et al., 2019a).

### 2.7. Extraction of transcription factors modulate human miRNA expression

We have downloaded the experimentally validated TFs which bind to miRNA promoters and module it. We have considered those TFs that are expressed itself and that can ‘activate’ or ‘regulated’ miRNAs.

### 2.8. Identification of the host epigenetic factors genes

We used EpiFactors database (Medvedeva et al., 2015) to find human genes related to epigenetic activity.

### 2.9. Mapping of the human proteins in cellular pathways

We have utilized KEGG mapper tool (Kanehisa and Sato, 2020) for the mapping of deregulated genes SARS-CoV-2 interacting host proteins in different cellular pathways. We then searched and targeted the pathways which are found to be enriched for SARS-CoV-2 deregulated genes. From these pathway information, we have manually sketched the pathways to provide a brief overview of the interplay between SARS-CoV-2 and host immune response, their outcomes in the viral pathogenesis.

## 3. Results

### 3.1. Differentially expressed genes found in SARS-CoV-2 infection are involved in different important cellular signaling pathways

We wanted to identify which pathways are to be modulated upon the infection of SARS-CoV-2 and their uniqueness from SARS-CoV infection. To find those, we have performed the enrichment analysis using the differentially expressed genes of both SARS-CoV and SARS-CoV-2 by Gitools (Perez-Llamas and Lopez-Bigas, 2011) using GOBP, GOMF, KEGG pathways, Bioplanet pathways, and Wikipathways modules.

We have identified 387 upregulated and 61 downregulated genes in SARS-CoV infection (analyzing GSE17400), and 464 upregulated and 222 downregulated genes in SARS-CoV-2 infection (analyzing GSE147507) (Supplementary file 2). Enrichment analysis of these differentially expressed genes showed that deregulated genes of SARS-CoV-2 infection can exert biological functions like-regulation of inflammatory response, negative regulation of type-I interferon, response to interferon-gamma, interferon-gamma mediated signaling, NIK/NF-kappaB signaling, regulation of apoptotic process, cellular response to hypoxia, angiogenesis, negative regulation of inflammatory response, zinc ion binding, calcium ion binding etc. which were not enriched for SARS-CoV infection (Figure 1A, 1B). Also, different organ specific functions like-heart development, kidney development etc. were only enriched for differentially expressed genes in SARS-CoV-2 infection (Figure 1A).

**Figure 1:**
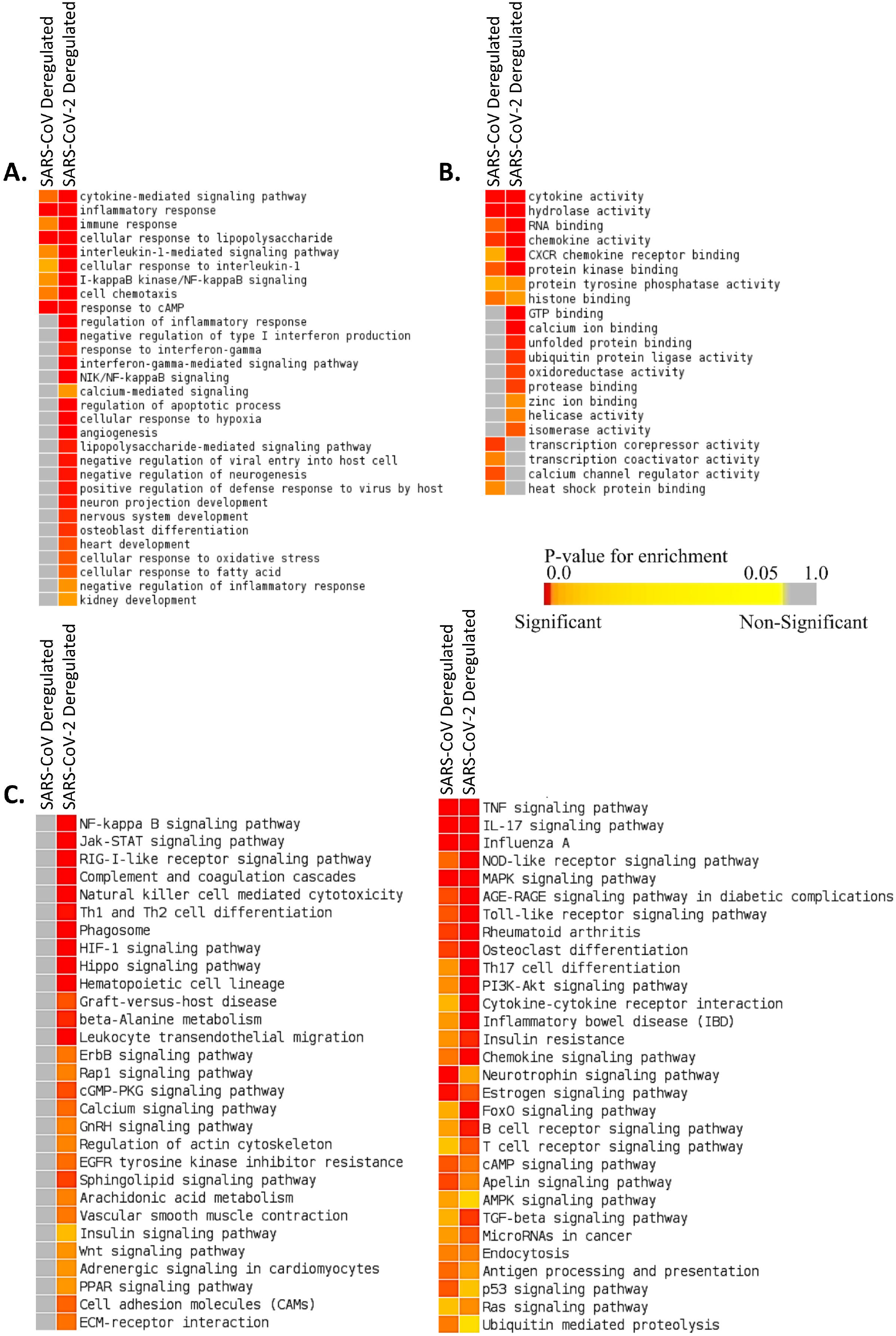
Enrichment analysis and comparison between deregulated genes in SARS-CoV and SARS-CoV-2 infections using **A.** GOBP module, **B.** GOMF module, **C.** KEGG pathway module. Significance of enrichment in terms of adjusted p-value (< 0.05) is represented in color coded P-value scale for all heatmaps. Color towards red indicates higher significance and color towards yellow indicates less significance, while grey means non-significant.

Deregulated genes of SARS-CoV-2 infection were found to be related to pathways like-NF-kappaB signaling, Jak-STAT signaling, RIG-I-like receptor signaling, Natural killer cell mediated cytotoxicity, Phagosome, HIF-1 signaling, Calcium signaling, GnRH signaling, Arachidonic acid metabolism, Insulin signaling, Adrenergic signaling in cardiomyocytes, PPAR signaling etc. (Figure 1C, Supplementary figure 1, 2) which were found to be absent for SARS-CoV infection.

### 3.2. Both SARS-CoV-2 interacting human proteins and deregulated genes from SARS-CoV-2 infection have immunological roles

To find out whether SARS-CoV-2 interacting human proteins and differentially expressed genes in SARS-CoV-2 infection are involved in same pathways, we have carried out an enrichment analysis using STRING (KEGG pathway module) (Szklarczyk et al., 2019) which is a functional protein-protein interaction network database.

From this analysis, we have found that both the deregulated genes of SARS-CoV-2 infection and the SARS-CoV-2 interacting human proteins are involved in several important immune signaling pathways, namely-IL-17 signaling, NF-kappaB signaling, TNF signaling, Toll-like receptor signaling, Phagosome, Apoptosis, Necroptosis, PI3K-Akt signaling, HIF-1 signaling, MAPK signaling etc (Figure 2). Also, signaling pathways like-Relaxin signaling, Rheumatoid arthritis, AGE-RAGE signaling pathway in diabetic complications etc. were also enriched (Figure 2).

**Figure 2:**
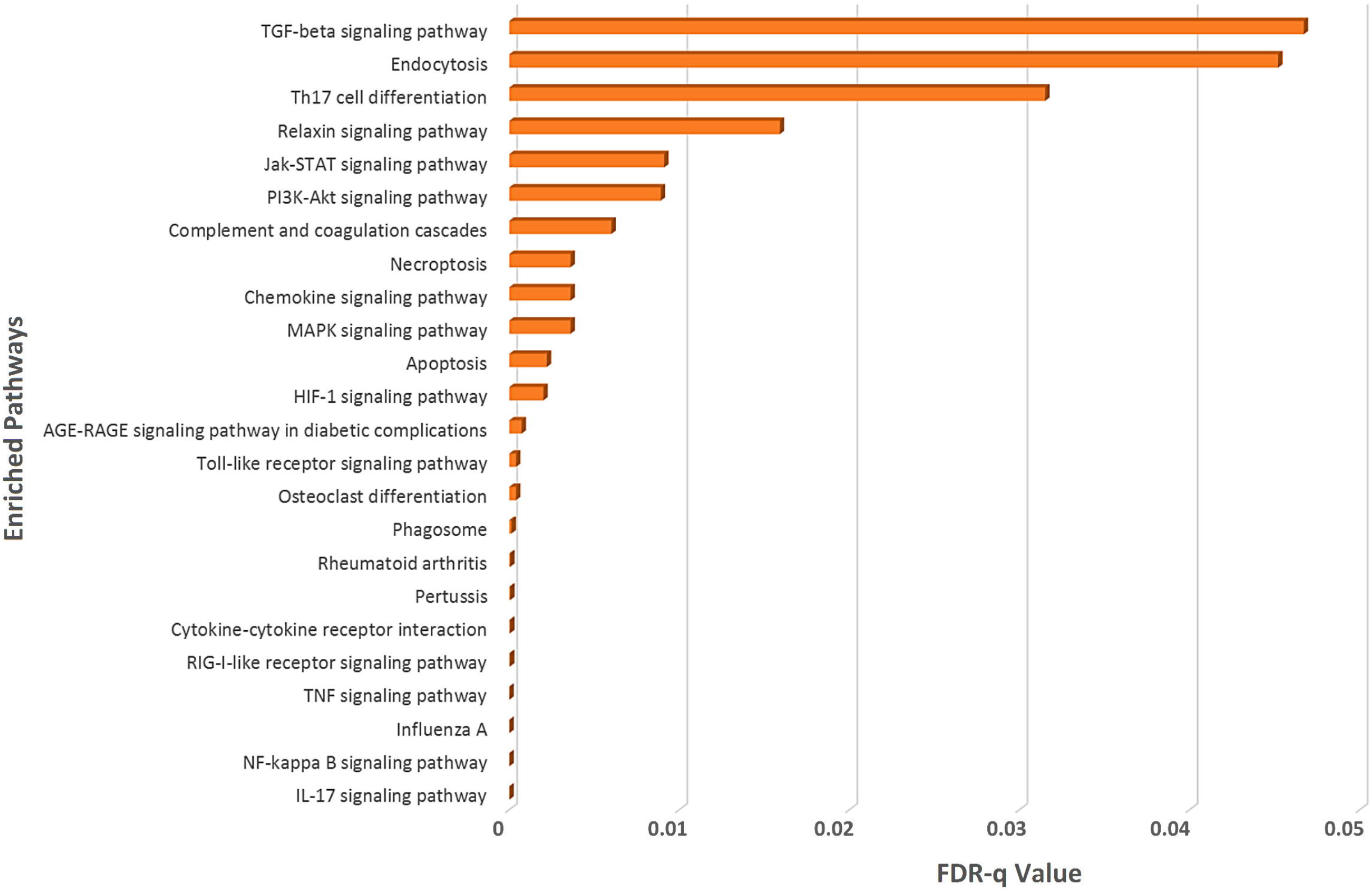
Enrichment analysis of deregulated genes in SARS-CoV-2 infections and human proteins interacting with SARS-CoV-2 proteins using KEGG pathway module of STRING database.

### 3.3. Differentially expressed genes in SARS-CoV-2 infection have different Transcription factor (TFs) binding preferences compared to SARS-CoV infection

We wanted to find out which transcription factors are preferentially binding to the promoters of the differentially expressed genes in SARS-CoV and SARS-CoV-2 infections and for this we have utilized CistromeDB (Zheng et al., 2018). We have identified 18 and 29 such TFs overrepresented around the differentially expressed genes of SARS-CoV and SARS-CoV-2, respectively (Figure 3, Supplementary figure 3). Among those TFs only 3 (NFKB1A, TNFAIP3, BCL3) were found to be common for deregulated genes of both infections. 19 of 29 TFs overrepresented for SARS-CoV-2 deregulated genes were also upregulated upon SARS-CoV-2 infections (Data not shown).

**Figure 3:**
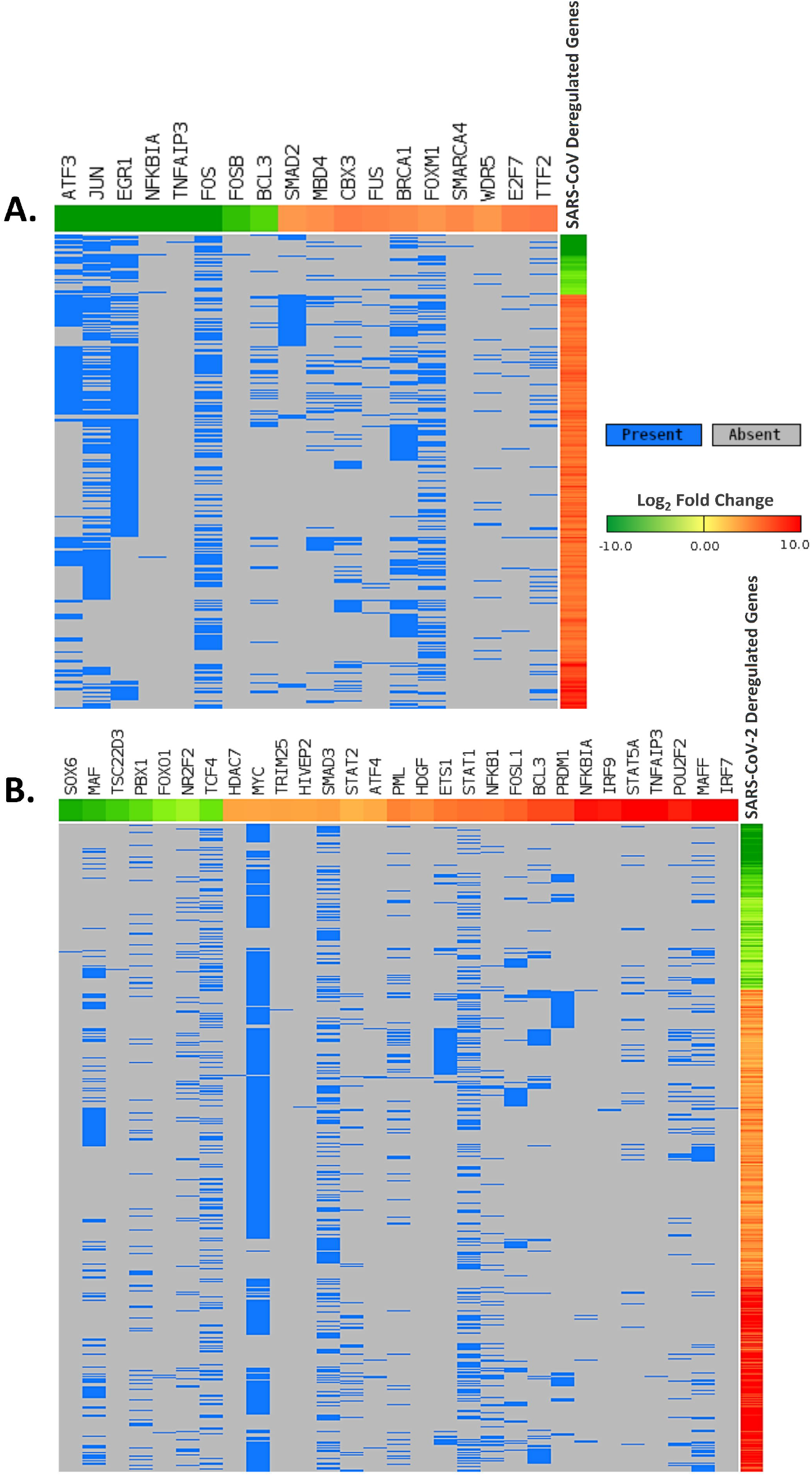
Differentially expressed genes in **A.** SARS-CoV and **B.** SARS-CoV-2 infections, and deregulated transcription factors which can bind around their promoters. Genes are represented vertically while the associated transcription factors are shown horizontally. Expression values of the TFs/genes are shown in Log2-fold change scale for both. Color towards red indicates upregulation and color towards green indicates downregulation. Blue color indicates presence and grey color indicates absence of a term.

### 3.4. Downregulated genes in SARS-CoV-2 infection can be targeted by different human miRNAs

Viral infection leading to the expression of different host miRNAs is a common phenomenon (Bruscella et al., 2017). To check if certain human miRNAs play a role behind the downregulation of the genes in SARS-CoV and SARS-CoV-2 infections, we have taken those miRNAs from miRTarBase database (Huang et al., 2019a) that can particularly target the downregulated genes in these infections. We have found 13 and 389 such candidate miRNAs targeting 17 downregulated genes in SARS-CoV infection and 123 SARS-CoV-2 infection, respectively (Supplementary file 3). Among those, only 7 miRNAs (hsa-miR-148a-3p, hsa-miR-146a-5p, hsa-miR-155-5p, hsa-miR-146b-5p, hsa-miR-27a-3p, hsa-miR-146b-3p, hsa-miR-141-5p) were found to be targeting in both infections.

### 3.5. Upregulated transcription factors in SARS-CoV-2 infection modulate different host miRNAs

To find out which differentially upregulated transcription factors can preferentially bind around the promoters of the host miRNAs and their associated functional roles, we have utilized TransmiR v2.0 database (Tong et al., 2019). We got 5 and 14 such upregulated TFs which might have a regulatory role on 14 and 90 host miRNAs in SARS-CoV and SARS-CoV-2 infections, respectively (Supplementary file 4). Though these transcription factors were completely different in both infections, we have found 6 host miRNAs (hsa-miR-146a, hsa-miR-146b, hsa-miR-155, hsa-miR-141, hsa-miR-200a, hsa-miR-27a) commonly modulated by TFs in both infections. Interestingly in SARS-CoV-2 infection, we have found 2 host miRNAs (hsa-miR-429 and hsa-miR-1286) which are influenced by upregulated transcription factor NFKB1 and have associations with several downregulated genes (BCL2L11, FKBP5 and TP53INP1 for hsa-miR-429; CLU for hsa-miR-1286).

### 3.6. Several host epigenetic factors can modulate the deregulation of gene expression in SARS-CoV-2 infection

Next we sought which epigenetic factors are themselves deregulated and if their deregulation could play a role in the overall differential gene expression. To do so, we have searched the EpiFactors database (Medvedeva et al., 2015) and identified 33 and 10 such epigenetic factors which were found to be deregulated in SARS-CoV and SARS-CoV-2 infections, respectively (Figure 4). Among the 10 factors found in SARS-CoV-2 infection, 6 (PRMT1, TRIM16, HDAC7, HDGF, DTX3L, PRDM1) were upregulated and 4 (PPARGC1A, PADI3, FOXO1, HELLS) were downregulated.

**Figure 4:**
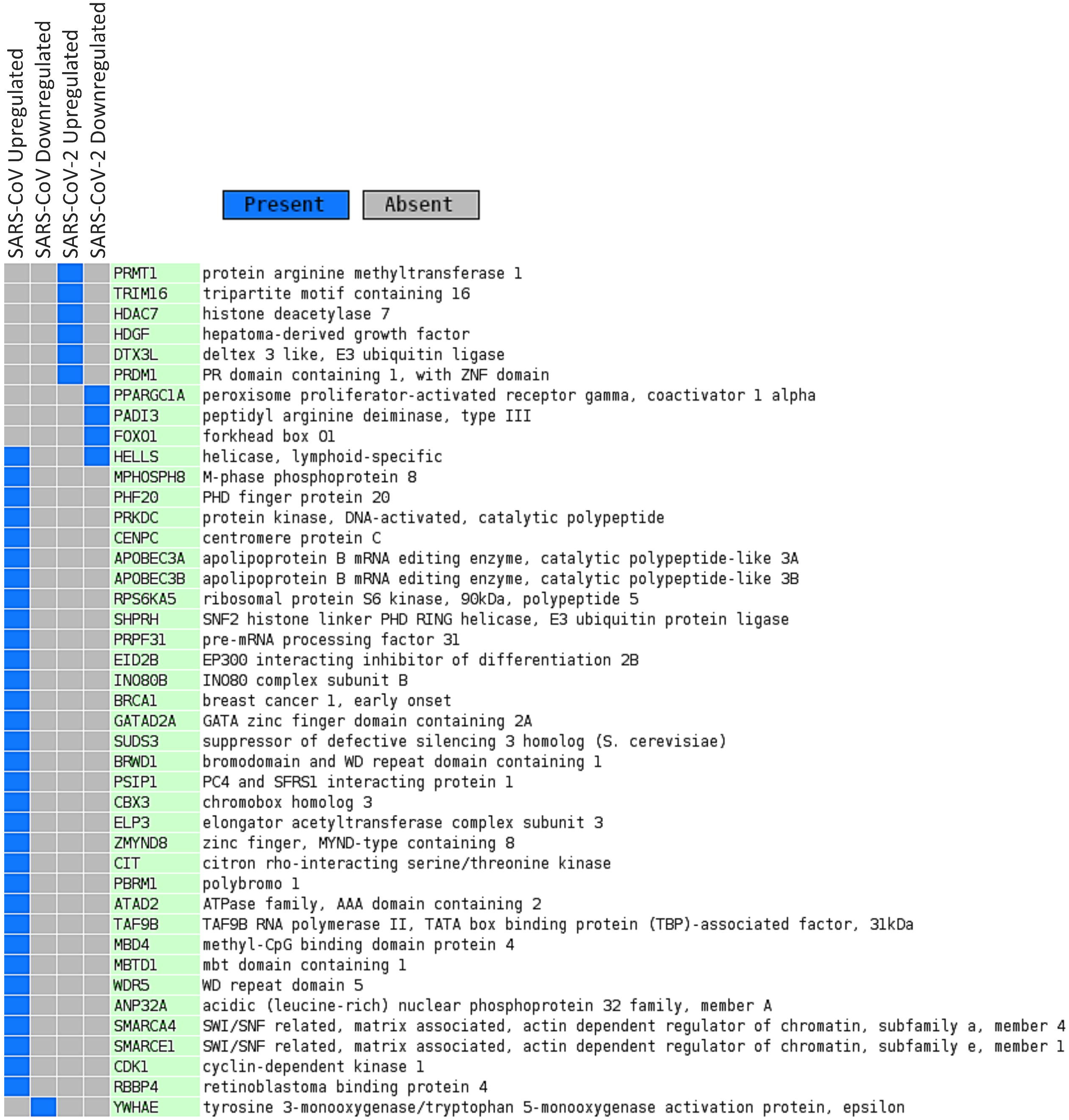
Deregulated epigenetic factors in SARS-CoV and SARS-CoV-2 infections. Blue color indicates presence and grey color indicates absence of a term.

### 3.7. Putative roles of viral proteins in immune evasion and pathophysiology of the COVID-19 are evident from various signaling pathways

Although there are some similarities between SARS-CoV and SARS-CoV-2 genetic architecture, it is yet to know if they modulate common host pathways. Also, it is largely unknown how SARS-CoV-2 uniquely exhibit some unique clinical features even having much similarities of the viral genes.

As now the probable genetic and epigenetic regulators behind the differential gene expression have been identified, we aimed to explore how these deregulated genes are playing a role in the battle between virus and host. To obtain a detailed idea of the outcomes resulting from viral-host interactions and how SARS-CoV-2 uses its proteins to evade host innate immune response, we have mapped the significantly deregulated genes and host interacting protein in different overrepresented functional pathways using KEGG mapper (Kanehisa and Sato, 2020). Analyzing the pathways, we have modeled several host-virus interactions in signaling pathways leading to the ultimate viral immune escape mechanisms. SARS-CoV-2 can blockade several signaling pathways like-HIF-1 signaling (Figure 5), Autophagy (Figure 6), RIG-I signaling (Figure 7), RIP1 mediated signaling (Figure 8), Beta adrenergic receptor signaling (Figure 9), Insulin signaling (Figure 10), Fatty acid oxidation and degradation pathway (Figure 10), IL-17 signaling (Figure 11), Toll-like receptor signaling (Figure 12), Phagosome (Figure 13A), Arachidonic acid metabolism (Figure 13B), PVR signaling (Figure 14) etc., aberration of these pathways might provide SARS-CoV-2 an edge over the host immune response. Also, SARS-CoV-2 can prevent the Relaxin downstream signaling (Figure 15) which plays a crucial role in lung’s overall functionality and its abnormal regulation might results in the respiratory complications found in COVID-19.

**Figure 5:**
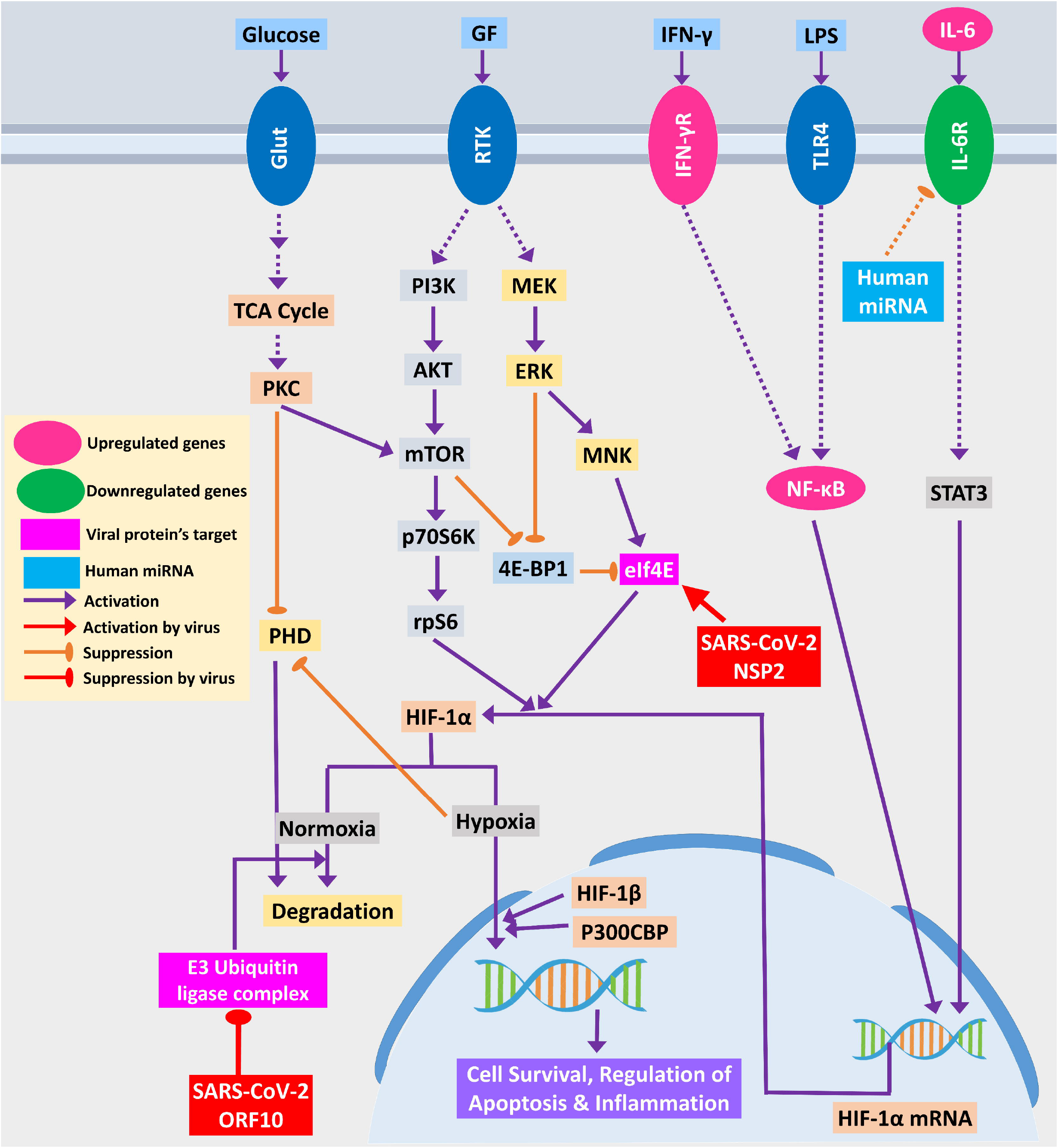
HIF-1 signaling pathway. Upregulated genes are in pink color while downregulated genes are in green color. Magenta and light blue representing viral protein’s target and host miRNAs, respectively. Violet pointed arrows indicating activation while orange blunt arrows indicating suppression. Red pointed and blunt arrows indicate activation and suppression by virus, respectively.

**Figure 6:**
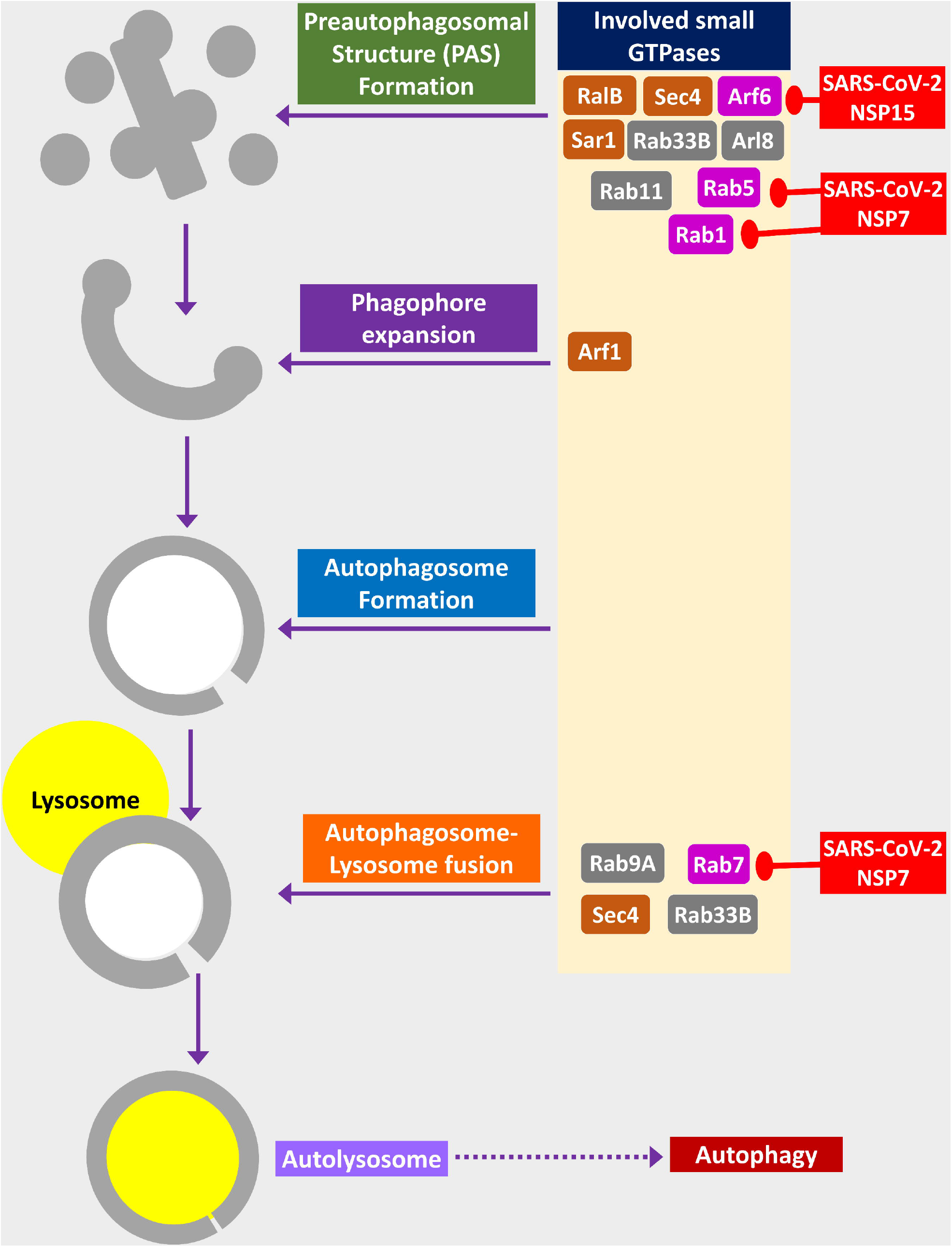
Small GTPases in Phagosome formation and maturation. Color codes are as in Figure 5.

**Figure 7:**
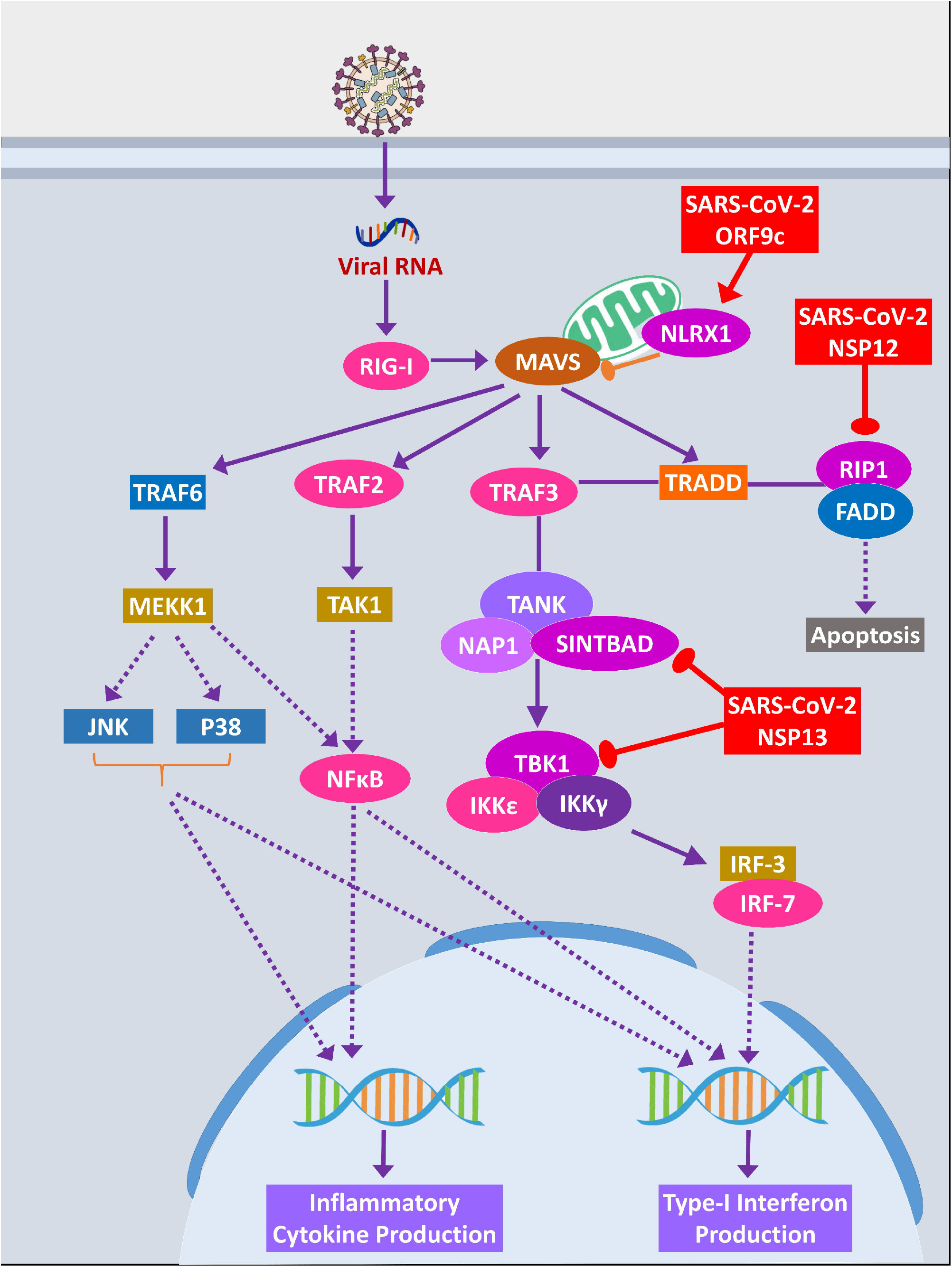
RIG-I signaling pathway. Color codes are as in Figure 5.

**Figure 8:**
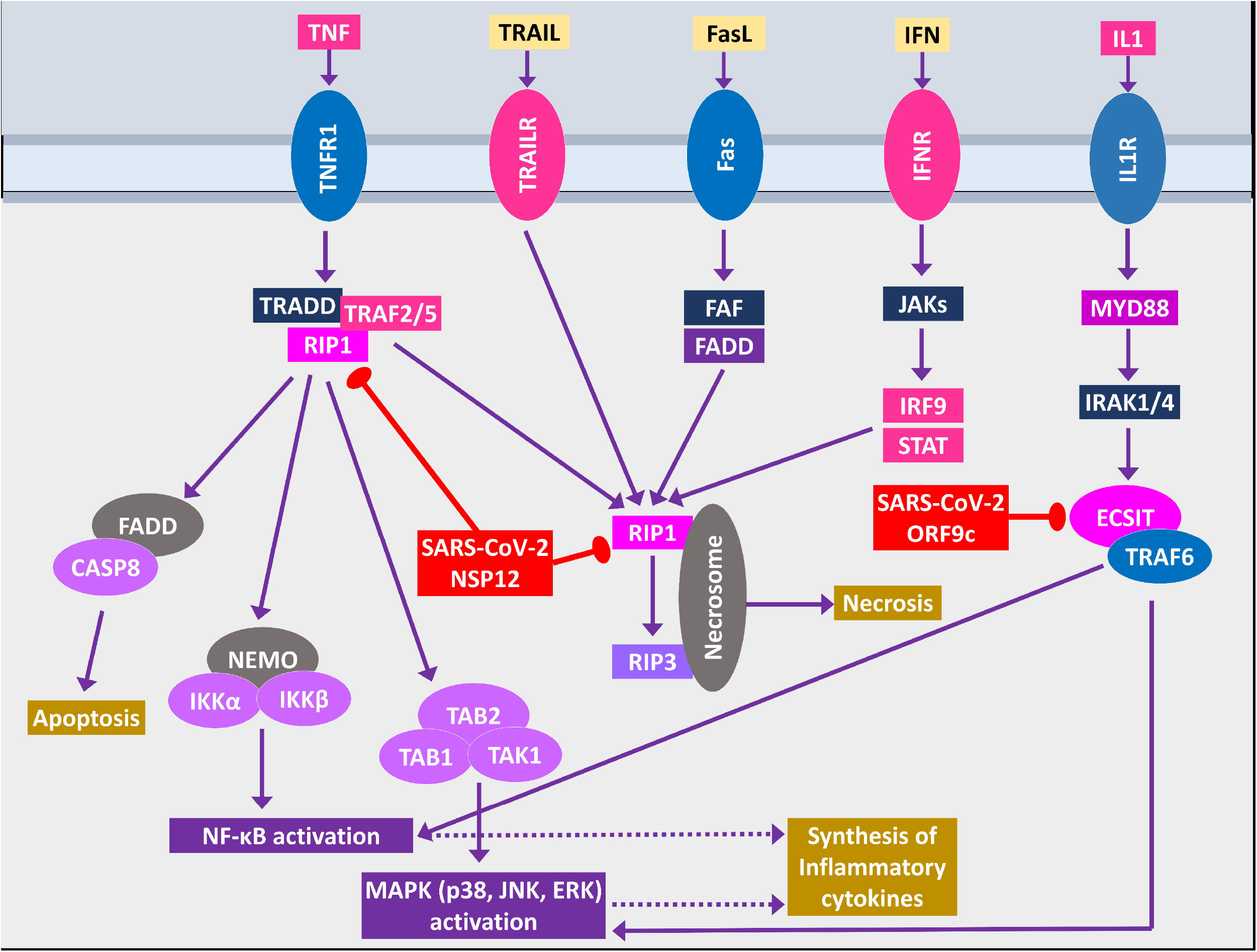
RIP1 and ECSIT signaling. Color codes are as in Figure 5.

**Figure 9:**
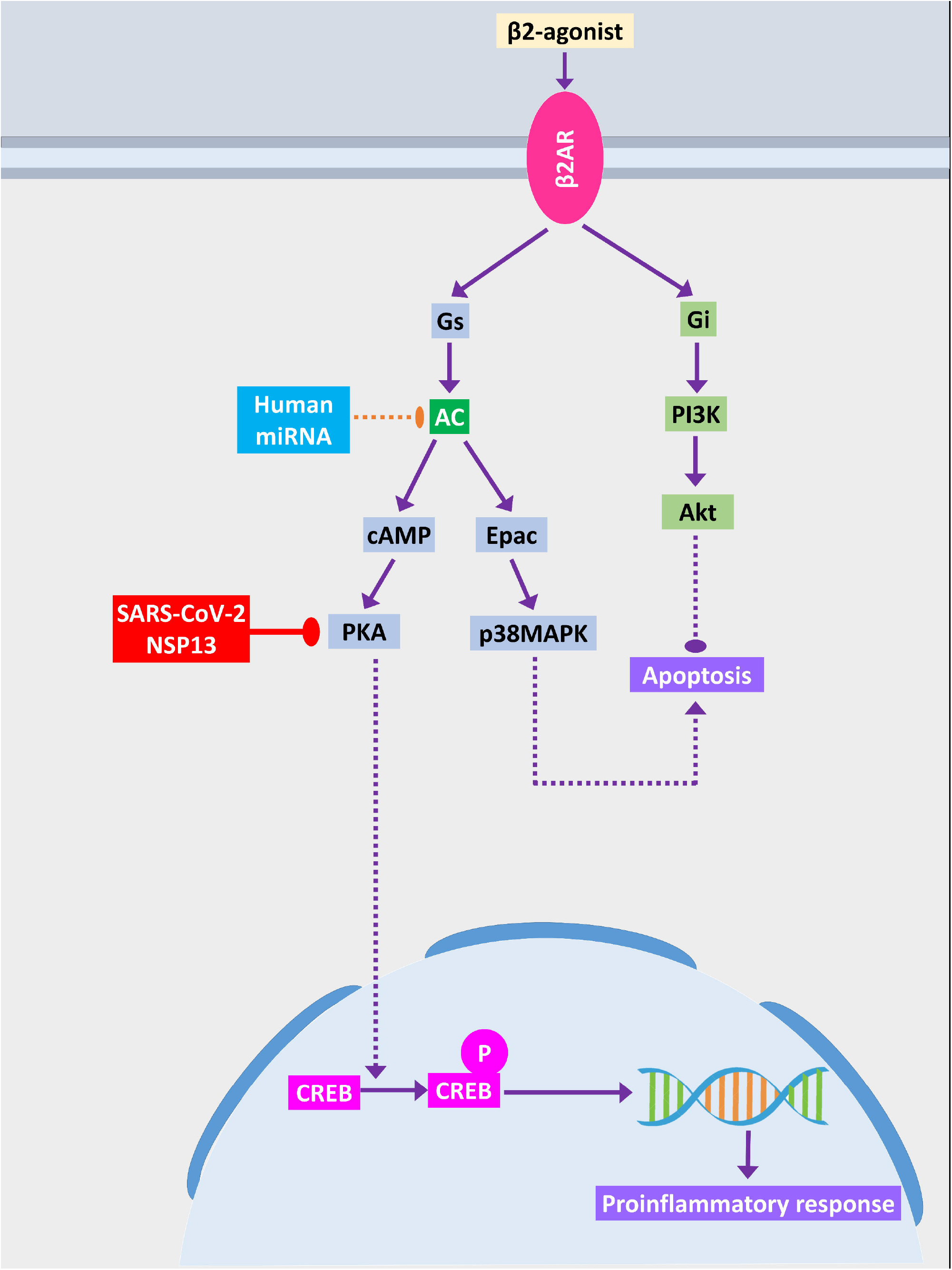
β2-adrenergic signaling. Color codes are as in Figure 5.

**Figure 10:**
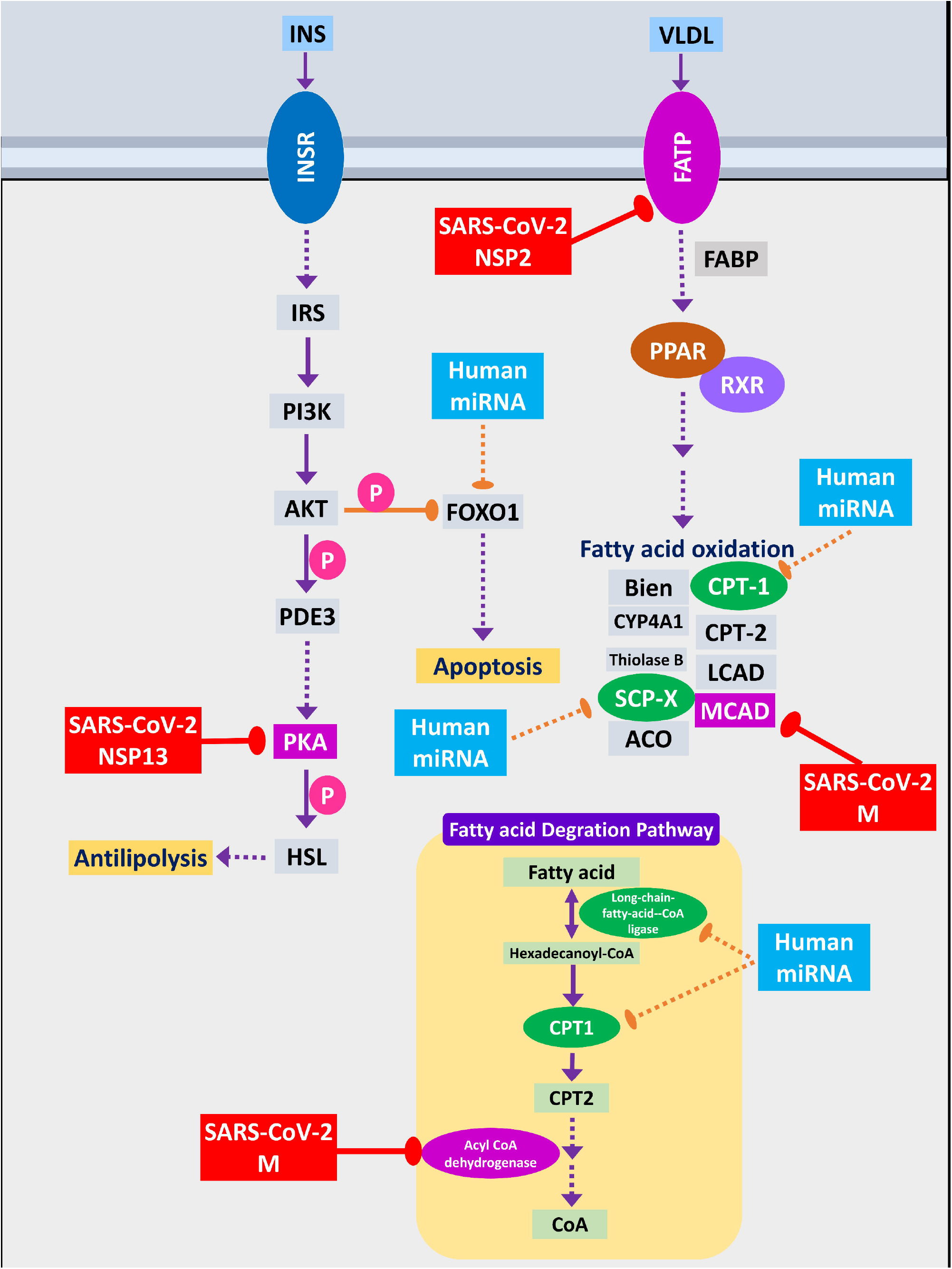
Fatty acid oxidation, fatty acid degradation and antilipolysis through insulin signaling. Color codes are as in Figure 5.

**Figure 11:**
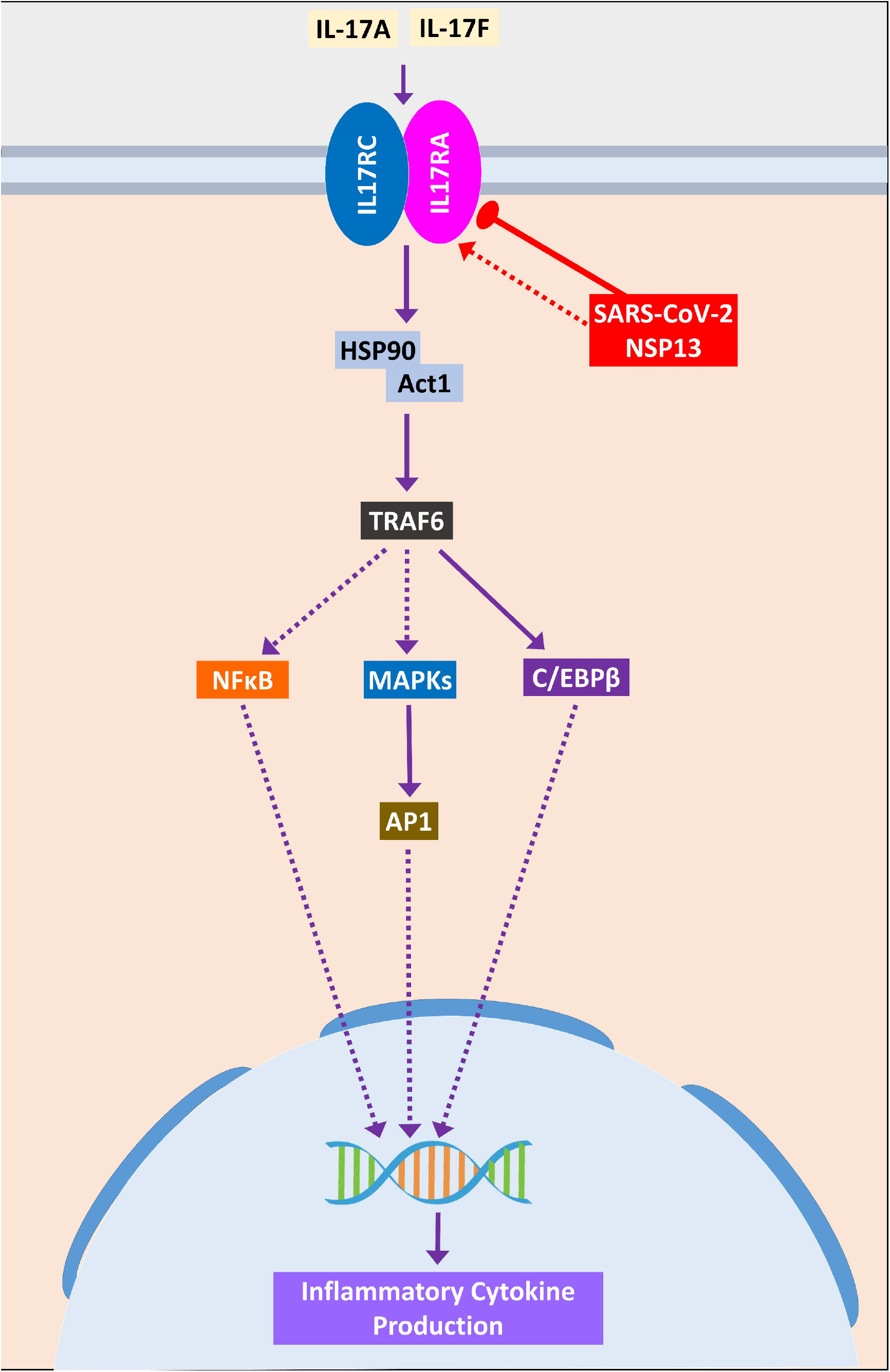
IL-17 signaling pathway. Color codes are as in Figure 5.

**Figure 12:**
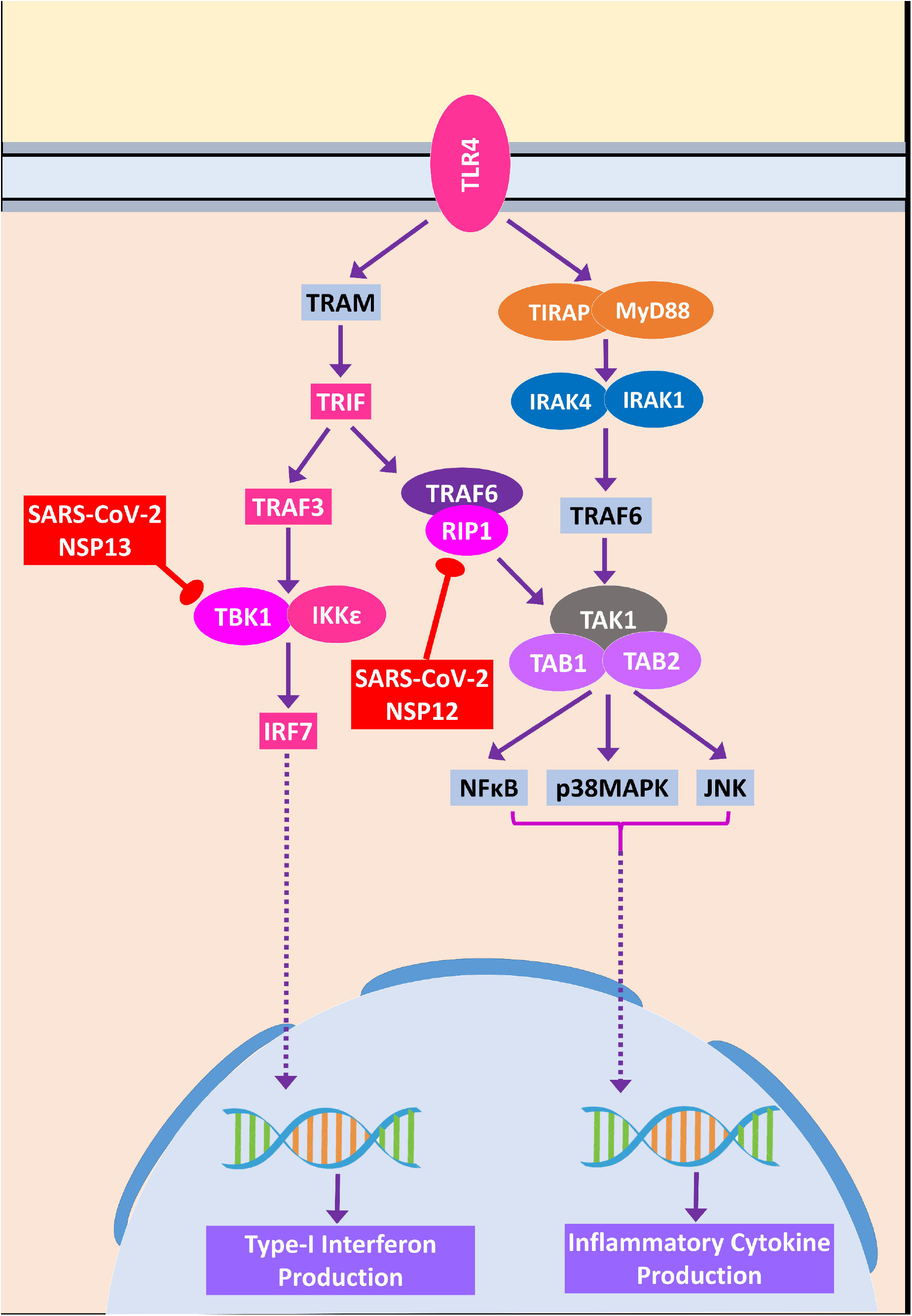
Toll-like receptor 4 signaling. Color codes are as in Figure 5.

**Figure 13:**
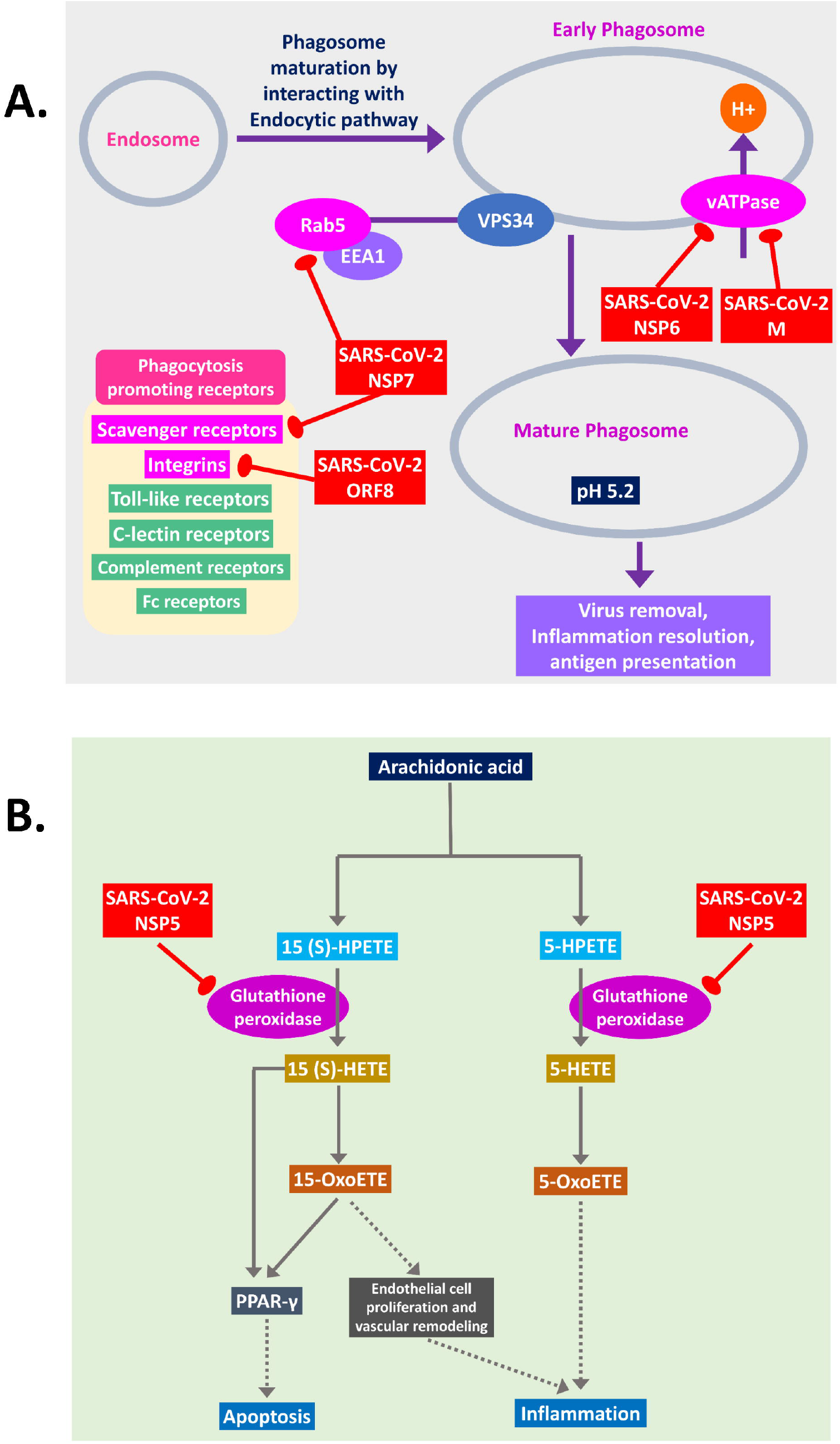
**A.** Phagosome maturation, and **B.** Arachidonic acid metabolism pathway. C Color codes are as in Figure 5.

**Figure 14:**
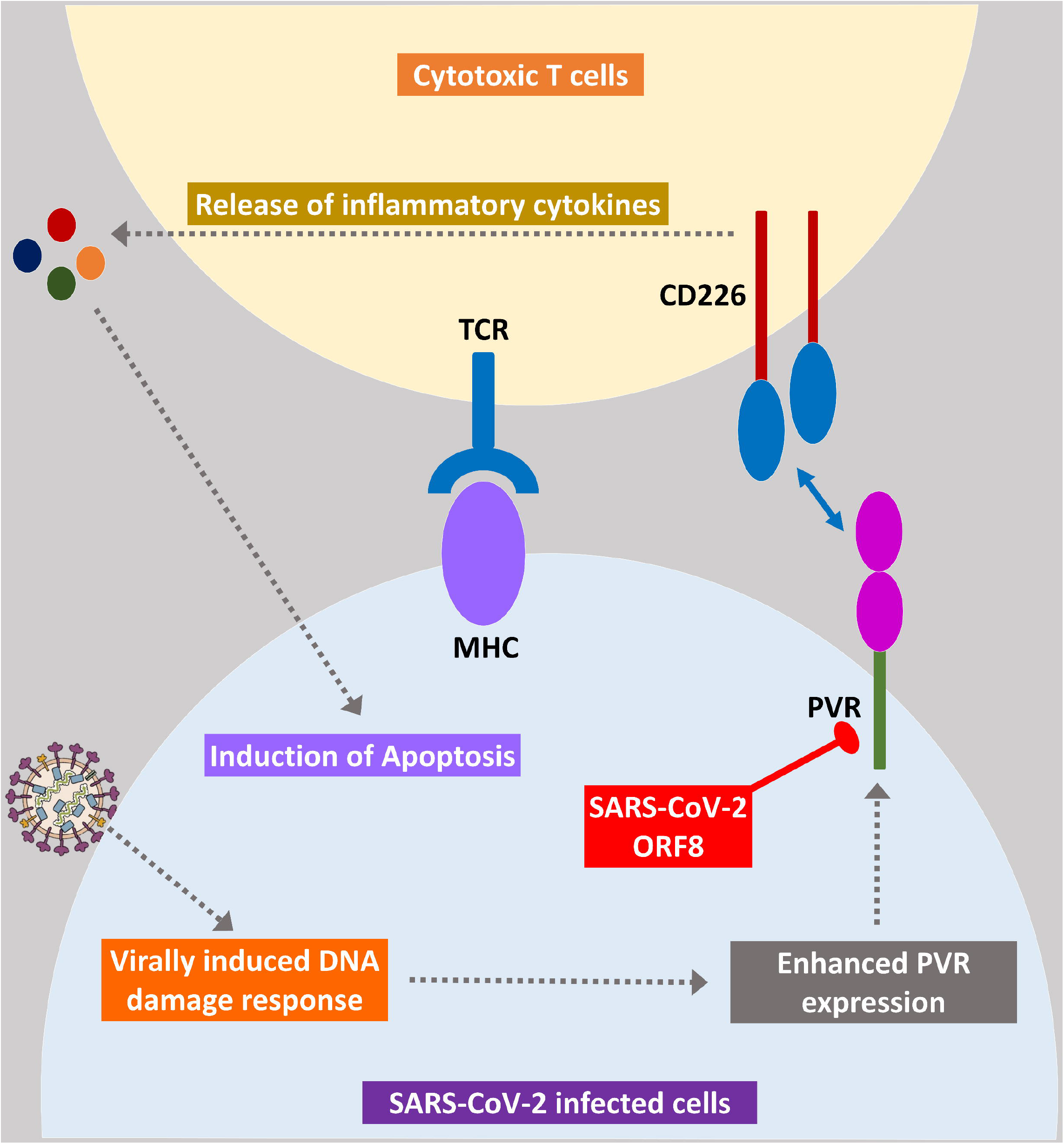
Cytotoxic T cells signaling using CD226 and PVR axis. Color codes are as in Figure 5.

**Figure 15:**
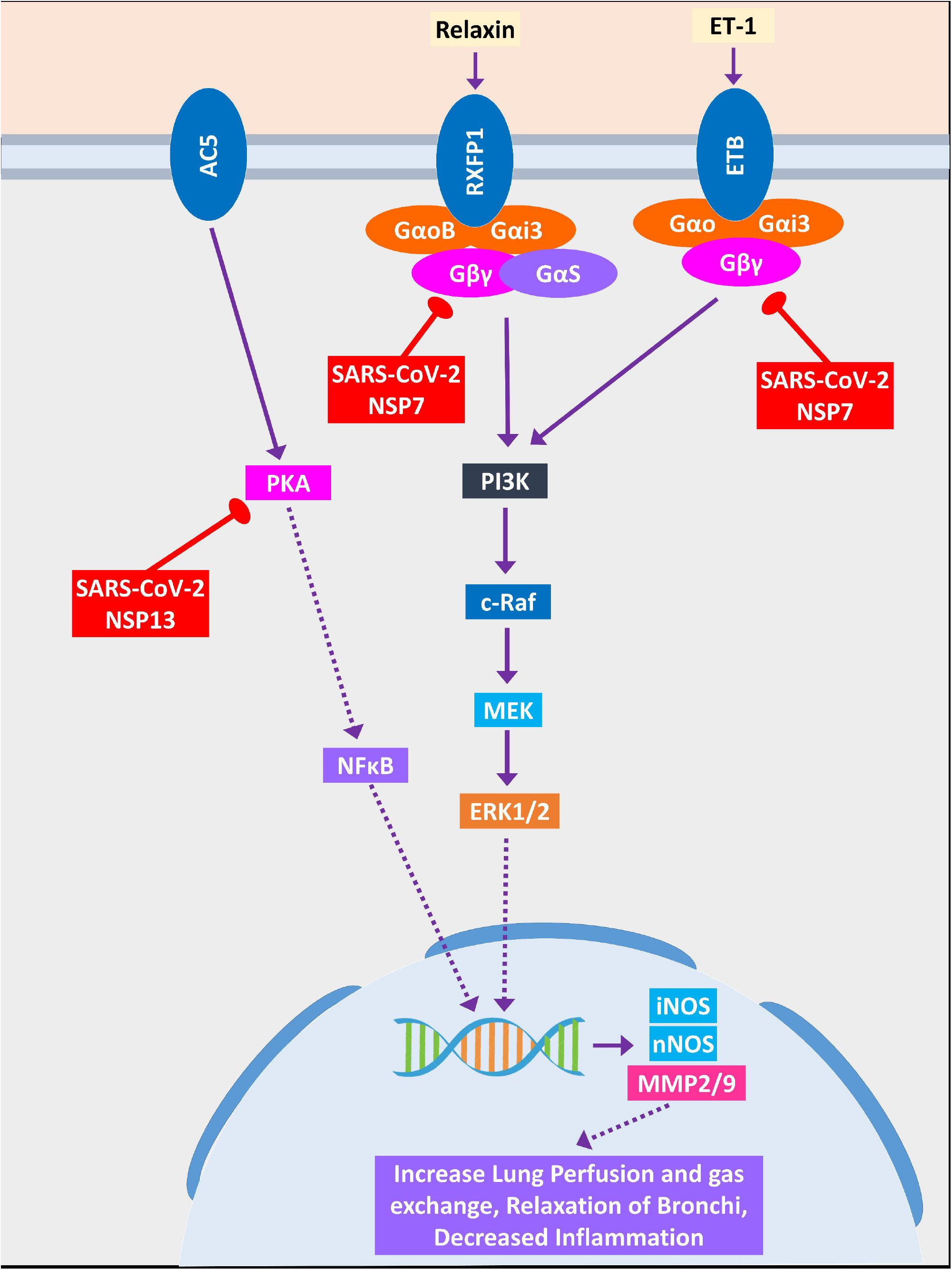
Relaxin signaling pathway in lungs. Color codes are as in Figure 5.

From previous studies we have compiled information on deregulated genes (Blanco-Melo et al., 2020) and virus-host interactome (Gordon et al., 2020) in SARS-CoV-2 infection, however, to get detailed pictures of the affected pathways, which is still remained obscure, we have investigated how our identified host genetic and epigenetic factors are playing a role and how viruses are utilizing those. Giving a closer look we have found some pathways which SARS-CoV-2 might be using but not SARS-CoV.

#### 3.7.1. SARS-CoV-2/host interactions lead to the observed lung related complications in COVID-19 patients

COVID-19 patients are reported to have suffer from hypoxic condition due to the breathing complications (Cascella et al., 2020). Hypoxia-Inducible Factor-1 (HIF-1) signaling pathway provides significant support mechanisms during this hypoxic condition by activating a wide ranges of other stress-coping mechanisms and ultimately leads to the survival of the stressed cells (Chen and Sang, 2016). So, if the infected cells utilize this survival mechanism, then the viruses propagating within these will also be saved. Thus, SARS-CoV-2 could be supporting this survival mechanisms as ORF10 protein binds and inhibits E3-ubiquitin ligase complex which degrades HIF-1α protein as many viruses modulates this complex for the benefits (Mahon et al., 2014) (Figure 5); it can also stimulate the functions of eIF4E (Eukaryotic Translation Initiation Factor 4E) for over production of HIF-1α as similar function was found for sapovirus protein VPg (Montero et al., 2015) (Figure 5).

SARS-CoV-2 mediated overexpression of HIF signaling products can lead to the pulmonary hypertension, acute lung injury etc. (Shimoda and Semenza, 2011) which might suggest the reason behind the frequent lung failure of critically infected patients. HIF signaling promotes hypoxia induced endothelial cell proliferations (Krock et al., 2011) which in turn might lead to aberrant clot formation in presence of amplified inflammation (Yau et al., 2015); which is frequently found in many COVID-19 patients.

Relaxin signaling plays significant role in maintaining lung’s overall functionality by maintaining lung perfusion and gas exchange, relaxation of bronchi and decreased inflammation in lungs (Alexiou et al., 2013). SARS-CoV-2 NSP7 protein can perturb this signaling by binding and inhibiting the relaxin receptors and prevents the productions NOS, MMP2/9 through PI3K to ERK1/2 axis (Figure 15). Another SARS-CoV-2 protein NSP13 might bind and block PKA, and its failure to activate NFκB may lead to the blockade of whole relaxin signaling (Figure 15). Aberration of this signaling pathway by SARS-CoV-2 possibly leads to the breathing complications in COVID-19 patients.

Binding of IL-17 receptor by SARS-CoV-2 NSP13 might increase the downstream signaling by activated TRAF6 to NFκB/MAPKs/CEBPB, which might cause some pathogenic inflammatory responses (Figure 11). Though IL-17 signaling is initially helpful in the host defense, still its aberrant expression might lead to pathogenic inflammatory responses leading to lung complications like-chronic obstructive pulmonary disease (COPD), lung fibrosis, pneumonia, acute lung injury etc (Gurczynski and Moore, 2018).

#### 3.7.2. SARS-CoV-2 proteins impede autophagy and phagocytosis to ensure its existence

During viral infection one important immune response is autophagy which destroys the virus infected cells and viruses are often found to modulate it for their survival (Ahmad et al., 2018). SARS-CoV-2 can inhibit the formation of autophagosome and phagosome using several of its proteins (Figure 6, 13A). SARS-CoV-2 NSP15 and NSP7 might bind and inhibit the some small members of Rab, Arf GTPases family (Figure 6) necessary for autophagosome formation (Bento et al., 2013). SARS-CoV-2 NSP7 and ORF8 can inhibit several phagocytosis promoting receptors like-scavenger receptors, integrins (Figure 13A); SARS-CoV-2 NSP6 and M protein might block vATPase, thus preventing the lowering of pH inside phagosome and preventing its maturation (Figure 13A); SARS-CoV-2 NSP7 prevents the phagosome-endosome/lysosome fusion by targeting Rab5 GTPase, which might results in virus pneumonia (Jakab et al., 1980) (Figure 13A).

#### 3.7.3. SARS-CoV-2 infection hinders host apoptotic responses necessary for their growth

Apoptosis is one important intracellular host immune response to reduce further spread of viruses from the infected cells (Barber, 2001). Several signaling pathways are involved to elicit this apoptotic response inside the infected cells which can be suppressed by the viral proteins, for example-NSP12 is found to target RIP1, thus it might fail to relay signaling to CASP8/FADD mediated apoptosis, and necrosis by RIP1/RIP3 complex (Figure 8); NSP5 might block glutathione peroxidase which is involved in 15(S)-HETE production and ultimately 15-oxoETE production, thus apoptotic induction by these metabolites through PPARγ signaling axis will not take place (Powell and Rokach, 2015) (Figure 13B); ORF8 can block PVR-CD226 signaling in cytotoxic T cell mediated apoptosis (Figure 14) as many viruses were reported to block PVR from expressing in the infected cell’s membranes (Cifaldi et al., 2019). Also, infection induced host miRNAs in β2-adrenergic signaling (Figure 9), insulin signaling (Figure 10) can block apoptosis of the infected cell.

#### 3.7.4. Host antiviral inflammatory cytokine and interferon production pathways are perturbed by SARS-CoV-2 proteins

Cytokine signaling pathways play a major role in suppressing the viral infections (Mogensen and Paludan, 2001). Similar inflammatory cytokine production pathways are also found in human coronaviral infections (Fung and Liu, 2019). We found that SARS-CoV-2 proteins are interacting with the members of these pathways and which might alter the signaling outcomes of these pathways to reduce the overall production of virus infection induced inflammatory cytokines.

RIG-I (Retinoic acid-Inducible Gene I) signaling plays important role in producing antiviral inflammatory cytokines and interferons, and induction of apoptosis (Chan and Gack, 2015). SARS-CoV-2 ORF9c protein can activate NLRX1 to degrade MAVS which results in failure of inflammatory cytokine production by NFkB via TRAF2/TAK1 or TRAF6/MEKK1 pathways (Figure 7). NLRX1 activity was previously found to be upregulated by HCV infection (Qin et al., 2017). By binding TBK1 and SINTBAD, SARS-CoV-2 NSP13 might inhibit the interferon production by IRF3/IRF7 stimulation (Figure 7).

Previous study suggest that RIP1 (Receptor-Interacting Protein Kinase 1) signaling plays key role in human coronavirus infection (Meessen-Pinard et al., 2017). RIP1 along with TRADD and TRAF2/5 activate NFκB and MAPKs which then induce the production inflammatory cytokines; this signaling axis might be blocked by SARS-CoV-2 NSP12 protein-RIP1 interaction, therefore, inhibiting the antiviral mechanisms exerted by this signaling pathway (Figure 8). SARS-CoV-2 ORF9c can also inhibit ECSIT/TRAF6 signaling axis which plays pivotal antiviral roles by activating NFκB and MAPKs signaling (Lei et al., 2015) (Figure 8). ß2-adrenergic signaling plays important antiviral roles in respiratory virus infections (Ağaç et al., 2018). SARS-CoV-2 NSP13 interacts with PKA, which might inhibit the activation of CREB by PKA for producing antiviral inflammatory responses (Figure 9).

Previous studies showed that Arachidonic acids suppress the replication of HCoV-229E and MERS-CoV (Yan et al., 2019). 15-oxoETE, an arachidonic acid metabolism product which promotes pulmonary artery endothelial cells proliferation during hypoxia (Ma et al., 2014) which in turn results in vascular remodeling and leakage of inflammatory cytokines (Powell and Rokach, 2015). 5-oxoETE another arachidonic acid metabolism product that can also induce inflammation (Powell and Rokach, 2015), production of both compounds might be hindered by SARS-CoV-2 NSP5 as it interacts with an upstream metabolic enzyme glutathione peroxidase (Figure 13B).

Previously it was reported that IL-17 signaling enhances antiviral immune responses (Ma et al., 2019). SARS-CoV-2 NSP13 can bind IL-17 receptor and inhibit the downstream signaling from IL-17 receptor to TRAF6 for activating NFκB/MAPKs/CEBPB signaling axis, thus decreasing the antiviral inflammatory responses (Figure 11).

During acute viral infections Toll-like receptor 4 (TLR4) signaling plays important roles in eliciting inflammatory responses (Olejnik et al., 2018). SARS-CoV-2 protein NSP13 interacts with TBK1 which might reduce the signaling from IRF7, resulting in less IFN-I productions; while NSP12 interacts with RIP1; as a result, the activation of downstream NFκB and MAPKs (p38 and JNK) pathways and induction of inflammatory responses from these pathways will be stalled (Figure 12).

#### 3.7.5. SARS-CoV-2 infection can negatively regulate fatty acid metabolism for its proliferation

Host lipid and fatty acid metabolism play crucial role in maintaining the viral life cycle and propagation inside the infected cells as viruses tends to utilize host metabolic pathways for its aids (Martín-Acebes et al., 2012). Cui et al. (2019) showed that impairment of fatty acid oxidation can lead to acute lung injury (Cui et al., 2019). SARS-CoV-2 NSP2 can interact with FATP receptor of fatty acid oxidation pathway, wheras M protein can interact and destabilize MCAD of fatty acid oxidation pathway; also, two other proteins were found to be induced human miRNA targets (Figure 10). MCAD deficiency leads to pulmonary haemorrhage and cardiac dysfunction in neonates (Maclean et al., 2005). So, this destabilization might lead to the acute lung injury during COVID-19. Increased fatty acid biosynthetic pathways are found in several viral infections for their efficient multiplications (Martín-Acebes et al., 2012), so it is logical for SARS-CoV-2 to inhibit the fatty acid degrading pathways. SARS-CoV-2 NSP13 protein were found to interact with insulin signaling mediated antilipolysis by PKA-HSL signaling axis; SARS-CoV-2 M and several host miRNAs can inhibit fatty acid degradation to CoA through CPT1-CPT2 metabolic axis (Figure 10), so that more fatty acids can be produced.

## 4. Discussion

The tug-of-war between the viral pathogens and infected host’s response upon the infection is a critical and complex relationship deciding the ultimate fate of an infection. Though most of the time successful removal of the virus is achieved through host immune response, still viruses have also evolved some immune evasion mechanisms to escape the immune surveillance of the host, thus, making the outcomes of the disease more complicated (Fung et al., 2020). Similar interactions were also found in other human coronaviruses which modulates the host immune responses (Fung and Liu, 2019; Fung et al., 2020). In this study, we have depicted how SARS-CoV-2 and host protein interactome leads to the probable immune escape mechanisms of this novel virus, along with the functional roles of other host epigenetic factors in this interaction as host epigenetic factors serve important roles in viral infections (Bussfeld et al., 1997; Paschos and Allday, 2010; Girardi et al., 2018).

Our analysis showed several transcription factors are capable of binding around the promoters of deregulated genes found in SARS-CoV-2 infection, which were absent in SARS-CoV infection. Some of these downregulated transcription factors in SARS-CoV-2 infection like-MAF, FOXO1 can elicit proviral responses (Kim and Seed, 2010; Lei et al., 2013). Also, some upregulated transcription factors in SARS-CoV-2 infection like-HDAC7 (Herbein and Wendling, 2010), STAT2 (Le-Trilling et al., 2018), ATF4 (Caselli et al., 2012), FOSL1 (Cai et al., 2017) can facilitate progression of viral life cycle and immune evasion in host. Though other upregulated transcription factors in SARS-CoV-2 infection TRIM25 (Martín-Vicente et al., 2017), SMAD3 (Qing et al., 2004), STAT1, IRF7, and IRF9 (Chiang and Liu, 2019) might play a role in the antiviral immunity upon infection.

Interestingly, we have found 2 miRNAs-hsa-miR-429 and hsa-miR-1286 whose associated transcription factors were upregulated and their target genes were downregulated, suggesting they might have some roles in this host-virus interactions. In RSV infection, hsa-miR-429 was found to be upregulated in severe disease conditions (Inchley et al., 2015). Other studies showed that this miRNA plays important role in promoting viral replication and reactivation from latency (Ellis-Connell et al., 2010; Bernier and Sagan, 2018). So, expression of this miRNA in SARS-CoV-2 infection can lead to similar disease outcomes.

Upregulated epigenetic factors in SARS-CoV-2 infections can be both a boon and a bane for the host, as factors like-TRIM16 (van Tol et al., 2017), DTX3L (Zhang et al., 2015) can provide antiviral responses; whereas factors like-PRDM1 (also known as BLIMP-1) (Lu et al., 2014), HDAC7 (Herbein and Wendling, 2010) can act as proviral factors.

Upregulated transcription factors like-SMAD3, MYC, NFKB1, STAT1 might be involved in the upregulation of hsa-miR-18a, hsa-miR-155, hsa-miR-210, hsa-miR-429, hsa-miR-433 which can in turn downregulate the HIF-1 production (Supplementary Figure 4) (Serocki et al., 2018). Also, epigenetic factors like PRMT1 can downregulate HIF-1 expression; while factors like HDAC7 can increase the transcription of HIF-1 (Luo and Wang, 2018). We found that when MYC, SMAD3, TNF transcription factors are upregulated, they can activate autophagy and apoptosis promoting miRNAs like-hsa-miR-17, hsa-miR-20, hsa-miR-106 (Supplementary Figure 4) (Xu et al., 2012). RIG-I signaling can induce miRNAs like-hsa-miR-24, hsa-miR-32, hsa-miR-125, hsa-miR-150 to neutralize viral threats (Li and Shi, 2013) and we also found these miRNAs can be transcribed by the upregulated TFs (Supplementary Figure 4). RIP1 can be targeted by induced hsa-miR-24 and hsa-miR-155 (Liu et al., 2011; Tan et al., 2018) in SARS-CoV-2 infection (Supplementary Figure 4). IL-17F can be targeted by upregulated hsa-miR-106a, hsa-miR-17, and hsa-miR-20a, while expression IL-17A can be modulated by upregulated hsa-miR-146a, and hsa-miR-30c (Supplementary Figure 4) (Mai et al., 2012). We also found that different upregulated TFs induced miRNAs like-hsa-miR-146a, hsa-miR-155 can downregulate TLR4 signaling (Supplementary Figure 4) (Yang et al., 2011).

From the enrichment analysis, we have found that deregulated genes were also involved in processes and functions like-heart development, kidney development, AGE-RAGE signaling pathway in diabetic complications, zinc ion binding, calcium ion binding etc (Figure 1A, 1B). Impairment of these organ specific functions might suggest the increased susceptibilities from COVID-19 patients having comorbidities. Zinc and calcium ions play significant roles in activating different immune response, aberrant regulation of these might be lethal for COVID-19 patients (Verma et al., 2011; Read et al., 2019).

Host antiviral immune responses encompassing different mechanisms like-autophagy, apoptosis, interferon signaling, inflammation etc. play fundamental role in neutralizing viral threats found not only in human coronaviruses (Fung and Liu, 2019) but also other other viral infections (Barber, 2001; Mogensen and Paludan, 2001; Ahmad et al., 2018). Viruses have also evolved hijacking mechanisms to bypass all those mechanisms for its survival (Fung et al., 2020). From our analyses, we have observed that SARS-CoV-2 might be playing similar mode of actions for its successful immune escape from the host immune surveillance. While all these mechanisms are supporting viral propagation, host suffers the adverse effects from these; resulting in the severe complications in COVID-19 patients.

All of these findings suggest that a very complex host-virus interaction takes place during the SARS-CoV-2 infections. During the infections while some host responses are playing significant impact in eradicating the viruses from the body, on the other hand virus modulates some critical host proteins and epigenetic machineries for its successful replication and evasion of host immune responses. Due to these complex cross-talk between the host and virus, the disease complications of COVID-19 might arise. Our study can be useful for designing further in depth experiments to understand the molecular mechanism of pathogenesis better and to develop some potential therapeutic approaches targeting these host-virus interactions.

## Author’s Contribution

ABMMKI conceived the project. ABMMKI and MAAKK designed the workflow. Both authors performed the analyses. MAAKK and ABMMKI wrote the manuscript. Both authors read and approved the final manuscript.

## Data Availability Statement

Publicly available data were utilized. Analyses generated data are deposited as supplementary files.

## Supplementary Figure legends

**Supplementary Figure 1:** Deregulated genes involved in significant pathways in SARS-CoV and SARS-CoV-2 infections. Blue color indicates presence and grey color indicates absence of a term.

**Supplementary Figure 2:** Enrichment analysis using deregulated genes in SARS-CoV and SARS-CoV-2 infections **A.** Bioplanet pathway module, **B.** Wikipathway module. Significance of enrichment in terms of adjusted p-value (< 0.05) is represented in color coded P-value scale for all heatmaps. Color towards red indicates higher significance and color towards yellow indicates less significance, while grey means non-significant.

**Supplementary Figure 3:** Detailed image of Figure 3. Genes are represented vertically while the associated transcription factors are shown horizontally. Expression values of the TFs/genes are shown in Log2-fold change scale for both. Color towards red indicates upregulation and color towards green indicates downregulation. Blue color indicates presence and grey color indicates absence of a term.

**Supplementary Figure 4:** Networks of **A.** SARS-CoV, and **B.** SARS-CoV-2 infection induced upregulated transcription factors and their trancribed miRNAs.

## List of supplementary files

**Supplementary file 1:** List of Human proteins and their associated interacting SARS-CoV-2 proteins.

**Supplementary file 2:** List of differentially expressed genes in SARS-CoV and SARS-CoV-2 infected cells.

**Supplementary file 3:** List of Human miRNAs targeting the donwregulated genes in SARS-CoV and SARS-CoV-2 infection.

**Supplementary file 4:** List of Upregulated Transcription factors in SARS-CoV-2 that can bind around the promoters of human miRNAs.

