## Supplementary figures and images for "SARS-CoV-2 proteins exploit host’s genetic and epigenetic mediators for the annexation of key host signaling pathways that confers its immune evasion and disease pathophysiology"

### Supplementary Figure 1

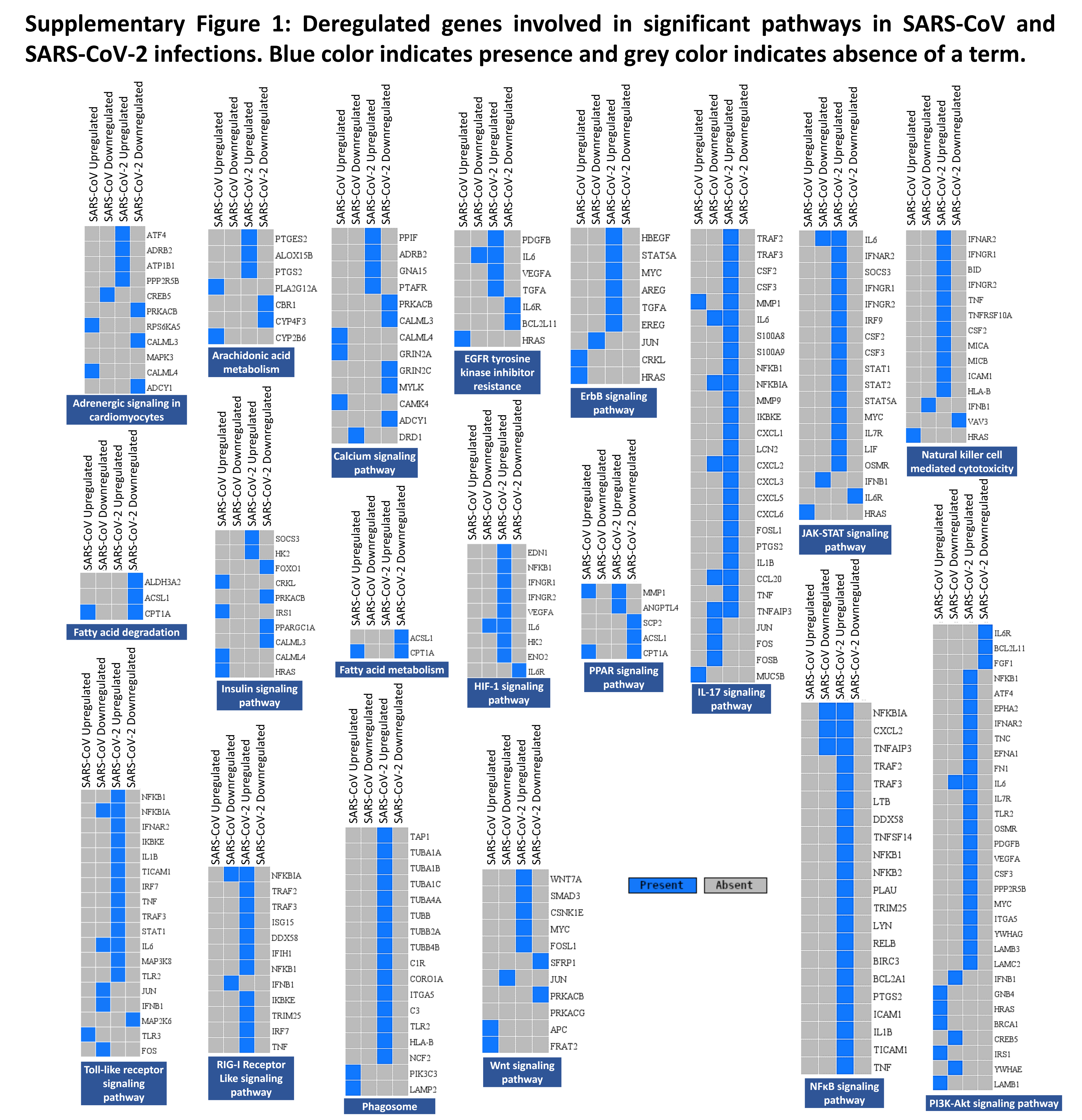

### Supplementary Figure 2

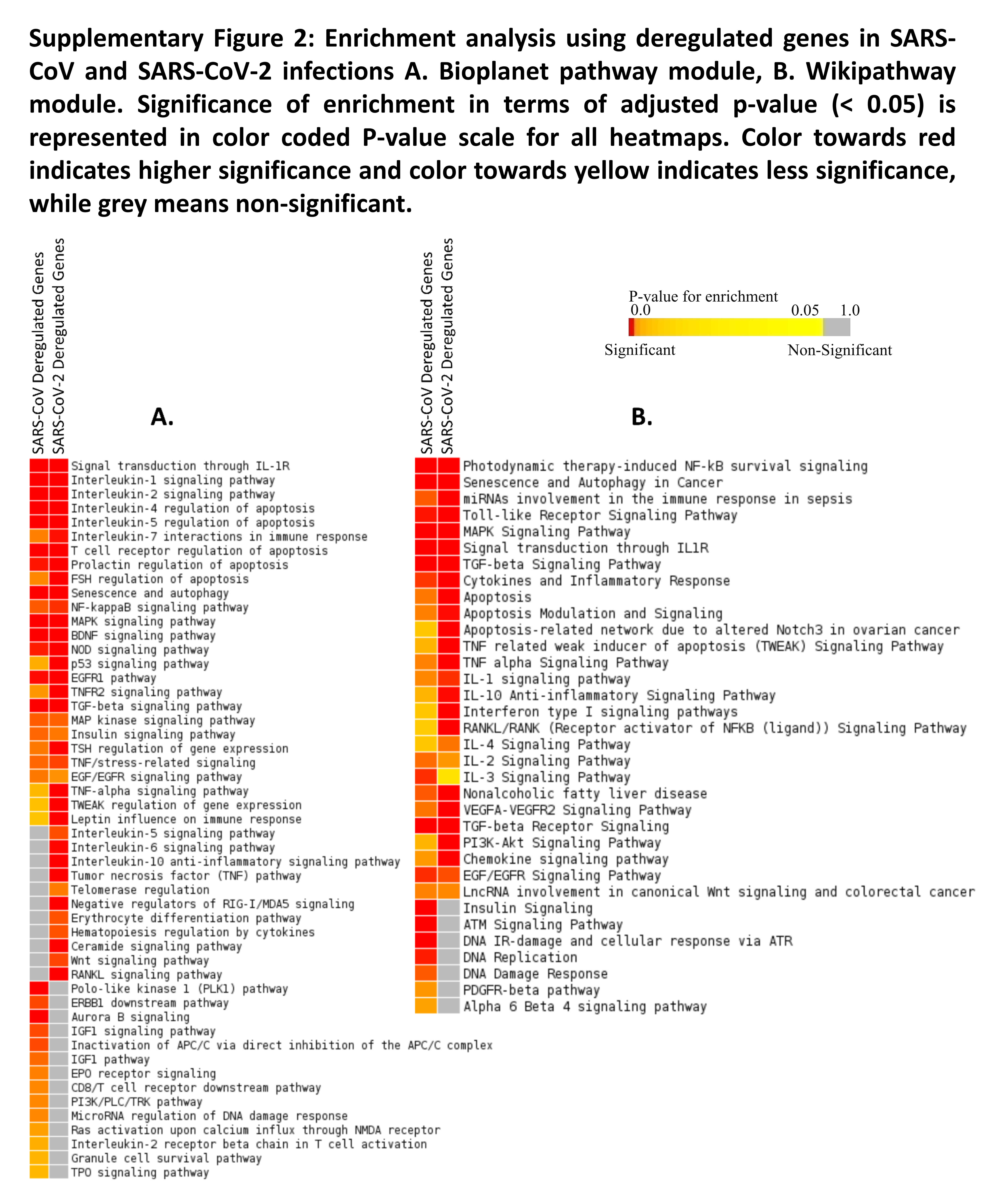

### Supplementary Figure 3

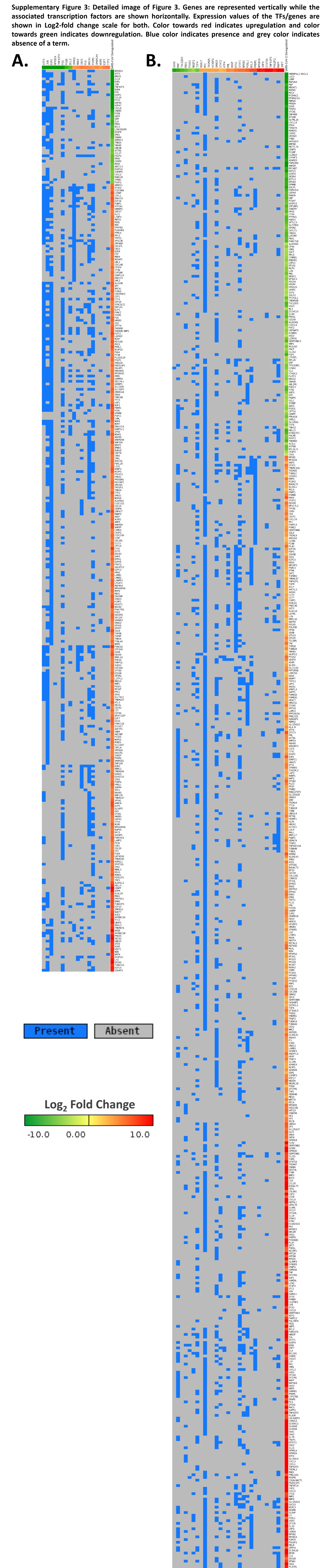

### Supplementary Figure 4

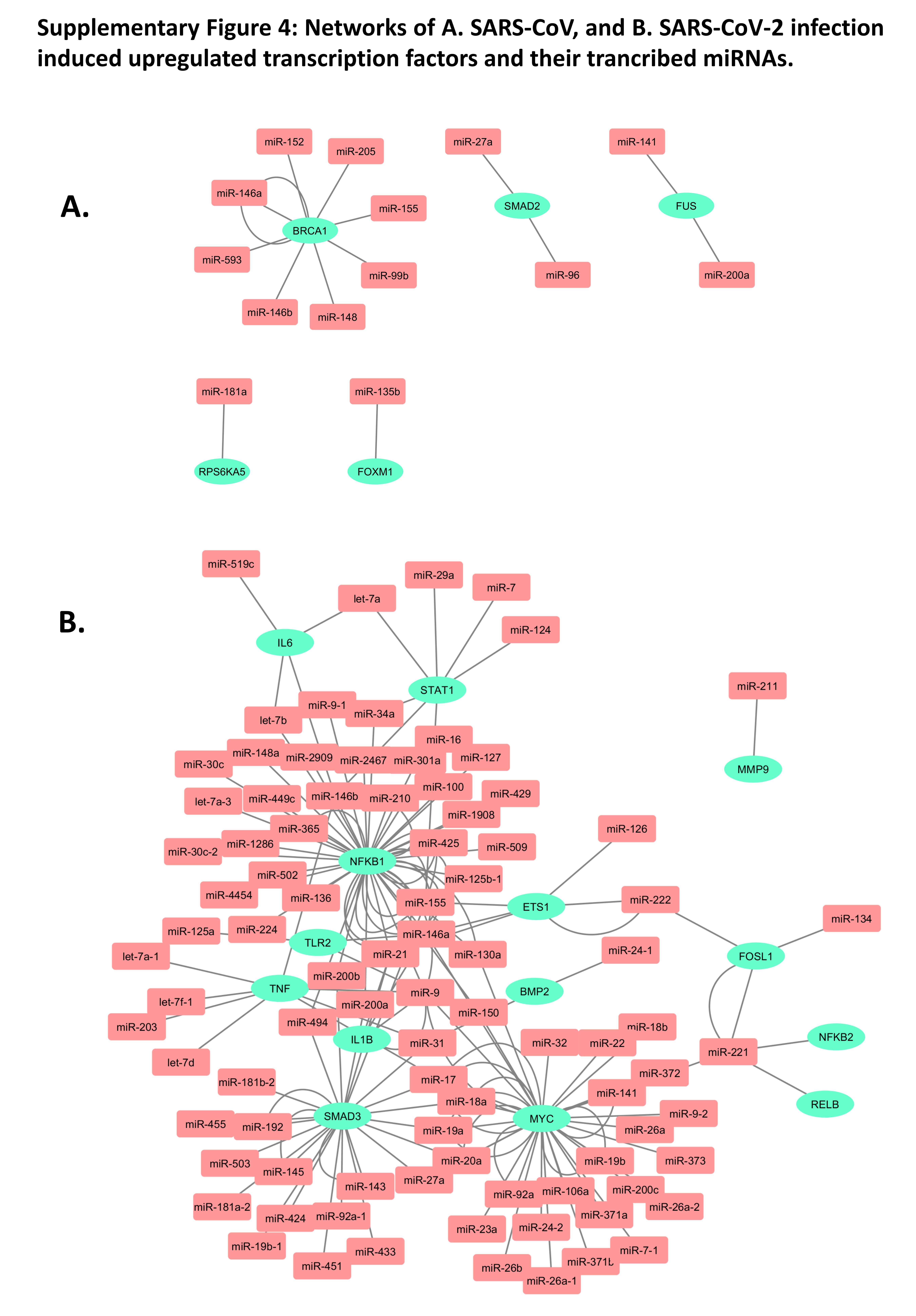
